# Identification of Juglone, a ‘first-in-class’ inhibitor of the human glutathione degrading enzyme, ChaC1, using yeast-based high throughput screens

**DOI:** 10.1101/2024.07.21.604522

**Authors:** Shradha Suyal, Chinmayee Choudhury, Deepinder Kaur, Anand K. Bachhawat

## Abstract

The cytosolic glutathione-degrading enzyme, ChaC1, is highly upregulated in several cancers, with the upregulation correlating to poor prognosis. The ability to inhibit ChaC1 thus becomes important in pathophysiological situations where elevated glutathione levels would be beneficial. As no inhibitors of ChaC1 are known, in this study we have focussed on this goal. We have initially taken a computational approach where a systemic structure-based virtual screening was performed. However, none of the predicted hits proved to be effective inhibitors. We also evaluated synthetic substrate analogs, but these too were not inhibitory. As both these approaches targeted the active site, we shifted to developing two high-throughput, robust, yeast-based assays that were active site independent. A small molecule compound library was screened using an automated liquid handling system using these screens. The hits were further analyzed using *in vitro* assays. Among them, juglone, a naturally occurring naphthoquinone, completely inhibited ChaC1 activity with an IC_50_ of 8.7 µM. It was also effective against the ChaC2 enzyme. Kinetic studies indicated that the inhibition was not competitive with the substrate. Juglone is known to form adducts with glutathione and is also known to selectively inhibit enzymes by covalently binding to their active site cysteine residues. However, juglone continued to inhibit a cysteine-free ChaC1 variant, indicating that it was acting through a novel mechanism. We evaluated different inhibitory mechanisms, and also analogues of juglone, and found plumbagin effective as an inhibitor. These compounds represent the ‘first-in-class’ inhibitors of the ChaC enzymes discovered using a robust yeast screen.

## Introduction

ChaC1 is a cytosolic glutathione-degrading enzyme found in higher eukaryotes including humans. It belongs to the γ-glutamylcyclotransferase family, and exclusively degrades reduced glutathione (and not oxidized glutathione) breaking it down into 5-oxoproline and cys-gly (1). ChaC1 also has a homolog, ChaC2 (66% similarity) which is found in all organisms from bacteria to man (2). ChaC2 also acts only on reduced glutathione but has far lower catalytic efficiency (20-fold lower), and is also expressed at lower levels (2,3). ChaC1, in contrast, is only present in higher eukaryotes and is expressed downstream of the Integrated stress response, under different stress and pathophysiological conditions. Owing to its exclusive activity towards reduced glutathione, ChaC1 upregulation results in not only glutathione depletion but in alterations in cellular redox ratios (4–7).

The upregulation of ChaC1 has been observed in many diseased conditions, cancers, and in ferroptosis (8). In cancer at least, this corresponds with increased proliferation and metastasis (9–11). These studies also indicate that ChaC1 expression in such diseased conditions is correlated to poor prognosis. Therefore, with such drastic consequences of ChaC1 induction, inhibiting the ChaC1 enzyme could be a potentially important intervention because it not only would result in higher glutathione levels in the cellular mileu, but also higher levels of the ratio of reduced to oxidized glutathione. However, no known inhibitors exist against either ChaC1 or ChaC2.

To increase glutathione levels in cells for therapeutic and other purposes, precursors (such as N-acetyl cysteine) and analogues of glutathione have been employed, since glutathione itself is not efficiently transported across the mammalian plasma membrane (12). A second approach has been to enhance glutathione biosynthesis by activating the transcription factor Nrf2 (13). A third approach that we propose can be considered, is to prevent glutathione degradation, by targeting the ChaC1 and ChaC2 enzymes.

There are several major hurdles towards identifying inhibitors of ChaC1. The first is that the Km of ChaC1 towards glutathione is in the range of 2 mM due to levels of glutathione in the cytoplasm ranging between 1-10 mM. Thus, finding efficient inhibitors that can outcompete the substrate could be potentially difficult. Secondly, there are no known inhibitors, so the search has to be begun *de novo*, and an inhibitor if found, would represent a ‘first-in-class’ inhibitor. Thirdly, the structure of ChaC1 is not known, and efforts to solve the structure have been hampered by the strong tendency of ChaC1 to aggregate *in vitro*. To address this last shortcoming, we recently modelled the structure of ChaC1 and also mapped the active site (14).

In this study, we have sought to identify inhibitors of ChaC1. Our initial approach was an *in silico* screening of large publicly available chemical libraries. However, the *in silico* screening of compound libraries was unable to provide us with a tentative lead, so we moved to an alternative strategy of screening a chemical compound library by developing two robust, yeast-based high-throughput screening assays. We expressed the ChaC1 enzyme from humans in two different strains of yeast showing opposite effects on their growth. In one assay, inhibition of ChaC1 prevents growth; in the other, its inhibition allows the growth of yeast cells. A small compound library of diverse structures was screened to identify the preliminary hits. These hits were further analyzed using *in vitro* assays, eventually leading us to a primary lead, juglone. Since juglone, is a naphthoquinone, with known mechanisms of action ascribed to it, we have explored the possible mechanisms by which juglone might be inhibiting ChaC1. Our results indicate that juglone, a ‘first-in-class’ inhibitor of ChaC1, inhibits ChaC1 and ChaC2 by mechanisms hitherto unknown for this naturally occurring metabolite.

## Experimental Procedures

### Virtual screening of compound libraries against the modelled ChaC1 structure

The virtual screening of the compound libraries involved three distinct steps which are outlined below.

### Generation of an interaction grid for molecular docking

The homology-modelled human ChaC1 structure was considered for virtual screening. Glutathione was docked to the model to generate a ChaC1-glutathione complex. Twenty snapshots were collected at every 10 ns interval from the MD trajectories of ChaC1. In each snapshot, the ligands were rescored in place and XP GScore and MMBGSA-binding energy (dG_bind) of glutathione with ChaC1 in each snapshot were calculated and averaged. The averaged docking score and MMGBSA-dG bind values were calculated to be -8.02 and -52.04 kcal/mol respectively, which were later considered while setting cut-off values for virtual screening. As all the receptor structures were prepared for docking before MD simulations, these snapshots were directly used for grid generation. In each snapshot, the glutathione conformation was rescored in place using Glide XP. “Receptor Grid Generation” module of Schrödinger was utilized to define interaction grids for molecular docking keeping glutathione as grid centers. The size of the interaction grid, was fixed to the default values 12Å for the inner box and 20Å as the outer box to facilitate a docking of slightly bigger molecules.

### Compound Dataset Preparation

A library containing a consolidated list of 5,30,881 drug like compounds from Asinex and 1,32,883 natural products from ZINC was prepared and used for virtual screening. These molecules were subjected to preparation in LigPrep (15) generating their ionization states at pH 7.0 (±2.0) using Epik ionizer, and three lowest energy conformers were retained for each compound. The prepared compound conformations were subjected to docking-based virtual screening.

### Virtual screening

The three best energy conformations of each compound were subjected to the virtual screening funnel. Initially, the Glide (16) module of the Schrödinger software package was used for performing high throughput virtual screening (HTVS), which employs a fast but less accurate scoring function. This was used to quickly eliminate the candidates that were unlikely to bind. This yielded 5000 compounds, which were further narrowed down to 1000 compounds, using the Standard Precision (SP) docking. Glide SP conducts a thorough sampling of the remaining ligands with a more accurate scoring function to refine the binding poses. Next, the screening process was made stringent downstream by employing a more computationally intensive method called Extra Precision (XP) docking, which facilitates the docking of compounds at a rate of approximately 2 minutes per molecule. Glide XP docking was used as the third filter where a Gscore cut-off of -8 was used to screen compounds with a lower value than the average docking score of glutathione. The threshold Glide XP score and MMGBSA binding affinity values were set at -8 and -60 Kcal/mol respectively (based on docking scores with the substrate, glutathione) for the final screening of compounds. All hits having GScore lower than -8 were subjected to MMGBSA-dG_bind calculation and hits with lower than -60 (a cut-off lower than the average MMGBSA-dG_bind of glutathione) were obtained for each grid. Now a consolidated list of 102 hits was prepared by combining and removing redundant hits obtained from all the 20 grids.

### Chemicals and Reagents

All chemicals used in the present study were of either analytical or molecular biology grades and were obtained from commercial sources. Media components were purchased from Difco, Merck, and HiMedia. Restriction enzymes and T4 DNA ligase were obtained from New England Biolabs. Phusion™ High-Fidelity DNA Polymerase was obtained from Thermo Scientific. Gel extraction kits and plasmid miniprep columns were obtained from Bioneer Inc. (Daejeon, South Korea) or Agilent Technologies. Oligonucleotides were purchased from Merck. Gossypin, Juglone, Catechin hydrate (Gossypin), Deoxycholic acid, Carbenoxolone disodium salt, S-methyl glutathione, oxidized glutathione, and Glutathione were purchased from Merck. Pro-GA was purchased from Funakoshi Co., Ltd.

Ten milligrams of each of the compounds ZINC12890221, ZINC13518810, ZINC1782149, ZINC1211353, ZINC857569, ZINC798255, ZINC9057554, ZINC20763370, and ZINC4940022 were procured from MolPort. 25 milligrams of each of the six glutathione analogs; γ-glu-leu-gly, γ-glu-met-gly, β-asp-cys-gly, β-asp-ser-gly, γ-glu-gly-gly, and γ-glu-ser-gly were custom synthesized from GenScript.

### Structures of compounds

The structures of virtual screening hits were downloaded from the ZINC15 database. Structures for juglone, p-benzoquinone, 2,5-dihydroxy 1,4 benzoquinone, DDQ (2,3-Dichloro-5,6-dicyano-1,4-benzoquinone), 1,4-naphthoquinone, lawsone, and plumbagin were downloaded from PubChem.

### Strains and growth conditions

*E. coli* DH5α and BL21 (DE3) pLysS were used as cloning host and expression host respectively. The *S. cerevisiae* ABC1723 (MATα; *his3Δ1 leu2Δ0 lys2Δ0 met15Δ0 ura3Δ0ecm38Δ::KanMX4 dug2-2*) and ABC1976 (MATα;*his3Δ leu2Δ 0 lys2Δ0 ura3Δ0gsh1Δ:: KanMX*) were used for creating the final strains used in the screen [ABC6293 (MATα; *his3Δ1 leu2Δ0 lys2Δ0 met15Δ0 ura3Δ0ecm38Δ::KanMX4dug2-2pdr5Δ::*LEU2) and ABC6303(MATα;*his3Δ leu2Δ 0 lys2Δ0 ura3Δ0gsh1Δ:: KanMX pdr5Δ::*LEU2)]. The yeast strains were maintained in YPD (yeast extract, peptone, and dextrose). For growth assays, Synthetic Defined (SD) minimal media containing YNB (yeast nitrogen base), ammonium sulfate, and dextrose supplemented with histidine, leucine, lysine, and methionine (80 mg/liter) were used. Glutathione was used at a concentration of 200 μM. Yeast transformations were carried out using the lithium acetate method.

### Cloning of ChaC1 constructs in pRS416 vectors

For the construction of ChaC1 under ADH1 and CYC1 promoters, the TEF-ChaC1 construct was digested with *XbaI* and *SacI* enzymes and replaced with ADH1 and CYC1 promoters from p416ADH and p416CYC plasmids respectively.

### Cloning of the cys-free ChaC1 into a bacterial expression vector

The ChaC1_C_S mutant was generated using the p416TEF-SSSUC2-HsChaC1(C>S,91,168,189,211,213) plasmid as a template (Amandeep Kaur, PhD Thesis, 2016 https://shodhganga.inflibnet.ac.in/handle/10603/471811), utilizing *NdeI* and *XhoI* restriction enzymes for cloning into the pET23a vector. Details of the primers used are in Table S1.

### Growth assays using dilution spotting

ChaC1 cloned under promoters of different strengths and their respective vector controls were transformed individually in *S. cerevisiae* ABC 1723 (sulfur auxotrophic) and ABC 1976 (glutathione deficient) strains. The transformants were selected against the Ura auxotrophic marker. For growth assays, the transformants were grown overnight in a minimal medium with amino acid supplements and methionine (ABC1723)/glutathione (ABC1976) as a sulfur source. They were thoroughly washed twice to remove the media components and reinoculated in a fresh medium containing amino acid supplements, but without any sulfur source, at an OD_600 nm_ of 0.20. The cells were sulfur-starved for 5–6 hours until the early exponential phase was reached. Later these cells were harvested, washed, and resuspended in water to an OD_600 nm_ of 0.2. These suspensions were serially diluted to 1:10, 1:100, and 1:1000. Of these cell resuspensions, 10 μl were spotted on minimal medium plates containing methionine or glutathione as sole sulfur sources. The plates were incubated at 30°C for 2–3 days, and then the images were taken using the Bio-Rad Gel Doc™ XR^+^ imaging system.

### Gene deletion in yeast Saccharomyces cerevisiae

A PCR-based gene disruption method (17) was employed in the yeast strains ABC 1723 and ABC 1976. A disruption cassette, containing the sequence of Leucine auxotrophic marker flanked at both sides with a short homology sequence of 41/42 bp at the 5’ and 3’ termini of the target gene (Pdr5), was amplified by PCR using oligonucleotide primers (Table S1). The amplified disruption cassette was transformed into the yeast cells and selected by leucine prototrophy. The disruptants were further confirmed by phenotype.

### Compound Library

The primary screen was performed using 2320 compounds from the MicroSource SPECTRUM Diversity library (Discovery Systems, Inc., Gaylordsville, CT, USA) (www.msdiscovery.com/spectrum), provided at 10 mM concentrations in DMSO solution in microplate format, at a density of 80 compounds per plate (rows 1 and 12 vacant). Each compound had a minimum of 95%purity. These compounds were diluted to 2 mM concentrations in 50% DMSO, aliquoted in 96 well microplates, and stored at -80° C. The library was a gift from Dr. Deepak Sharma, CSIR-ImTech Chandigarh, India. All compounds were added to the culture at a final concentration of 100 μM in the primary screens.

### High-throughput assay development, compound treatment, and liquid handling

10 ml pre-cultures were started from single colonies and incubated overnight. Cells were harvested by centrifugation washed twice in sterile water and re-suspended to pre-defined cell densities in a sulfur-starved media as mentioned and allowed to grow until they reached their exponential phase. These cells were re-inoculated in a minimal medium with 100 μM glutathione (ABC 6293) and no sulfur source (ABC 6303) to an initial OD_600 nm_ of 0.15. Using a robotic liquid handling system (Tecan Freedom EvoR), 200 μl of the cell culture was dispensed using the LIH arm, into six sterile 96 well plates sequentially, except for the first and last columns which were reserved for no compound controls. Next, 10 μl of each compound was added from the source plate to the culture plate using the MCA96 tips and mixed up and down into each well, leading to a final concentration of 100 μM in the culture. No compound and only vector control cells containing 2.5% DMSO were added manually in the empty wells of the first column. The plates were re-lidded and incubated in a shaker maintained at 30℃. OD_600 nm_ was measured at an interval every 6 hours until 36 hours using a microplate reader (Tecan Infinite M200 monochromator). The shaker and microplate reader were integrated with the Liquid Handling system and the transfer of plates between the shaker and the reader was automated. Z’ prime and 3 standard deviations (SD) values were calculated using control values from 5 independent assay plates.

### Growth-optimized conditions for the screens on the robotics system

After several rounds of standardizations, the assay conditions were established as follows. The indicator cells were cultivated overnight with methionine (in the sulfur auxotroph) and GSH (in the *gsh1Δ* strain) in the pre-cultures and starved without these sources for 5-6 hours before re-inoculating at an initial OD_600 nm_ of 0.15 in media containing 100 µM of GSH (for the sulfur auxotrophic strain) and no GSH for the glutathione deficient strain. The cells were grown for 36 hours on continuous shaking at 30 °C in a plate reader with OD_600 nm_ measurement every hour.

### Expression and purification of recombinant human ChaC1, ChaC2 and yeast Dug1 proteins

C-terminal 8x His-tagged human ChaC1, ChaC2, and yeast Dug1 constructs cloned in pET23a vector were transformed in *E. coli* BL21 (DE3) pLysS. For expression of the proteins, the transformants were grown overnight in LB media containing ampicillin (100 μg/ml) and chloramphenicol (35 μg/ml). The secondary culture was re-inoculated at OD_600 nm_= 0.05 and allowed to grow until OD_600 nm_ = 0.5–0.6. The culture was induced with 0.5 mM IPTG and incubated at 37°C for 3 h. Cells were harvested and the cell pellet was lysed by sonication in a buffer containing 50 mM Tris pH 8.0, 300 mM NaCl, and 1 mM PMSF (Buffer A). The cell lysate was centrifuged at 15000 X g for 30 min at 4°C and the supernatant was incubated with Ni-NTA Agarose beads for 1 hour at RT. It was loaded onto a polypropylene column, and pre-equilibrated with Buffer A. The column was washed four times with Buffer A containing 50 mM imidazole. The bound protein was eluted in Buffer A containing 250 mM imidazole and dialyzed overnight. The purified protein was run on SDS-PAGE and detected using a coomassie stain. The purified protein was flash-frozen using liquid nitrogen and stored at -80℃ until further use.

### ChaC1p enzymatic assay for inhibition studies

5 ng of recombinantly purified human ChaC1 protein was pre-incubated with the 100 µM inhibitor compound. All compounds were dissolved in dimethyl sulfoxide (DMSO) and diluted with sterilized distilled water until the concentration of DMSO was 1% for 30 min in a 50 μl reaction mixture containing 50 mM Tris-Cl (pH 8.0) and 5 mM DTT. 2 mM of the substrate, glutathione, was immediately added to this mixture and further incubated for 30 min. The enzyme was inactivated by heating at 95℃ for 5 min. To this, 10 μl of reaction mixture containing 5 μg of Dug1p and 20 µM MnCl_2_ was added and incubated at 37℃ for another 1 hour. Cysteine liberated from the above reaction was measured using a ninhydrin-based method (18).

### ChaC2 assay

The ChaC2 assays were carried out similar to the ChaC1 assays. However, as ChaC2 has a much lower activity to compare the enzymes accurately, 1 μg of both ChaC1 and ChaC2 enzymes were incubated for 1.5 hours with varying concentrations of juglone.

### Dug1p assay

Dug1p was pre-incubated with varying concentrations of juglone for 30 min and later the substrate, cys-gly dipeptide (2.5 mM) was added to the reaction mixture and incubated for an hour. The cysteine liberated was measured using the ninhydrin-based method.

### *In silico* modelling and molecular dynamic simulations of juglone and juglone conjugates with ChaC1

Four model systems were constructed considered for molecular dynamics studies in order to explore the mechanism of inhibitory action of juglone. The ChaC1-Glutathione complex generated in our previous work (14) was used to build four different model systems as follows. First, the ChaC1-juglone complex was modelled by docking juglone at the glutathione binding site. Second, the ChaC1-glutathione-juglone complex was generated by further docking juglone to the ChaC1-glutathione complex. The third (ChaC1-juglone-C2-glutathione-adduct) and fourth (ChaC1-juglone-C3-glutathione-adduct) model systems were generated by docking two types of juglone-glutathione adducts at the glutathione binding site of ChaC1. The two adducts were modelled by attaching glutathione (–SH) with the 2^nd^ and 3^rd^ carbon atom of juglone as described in Scheme 1.

**Scheme 1.**
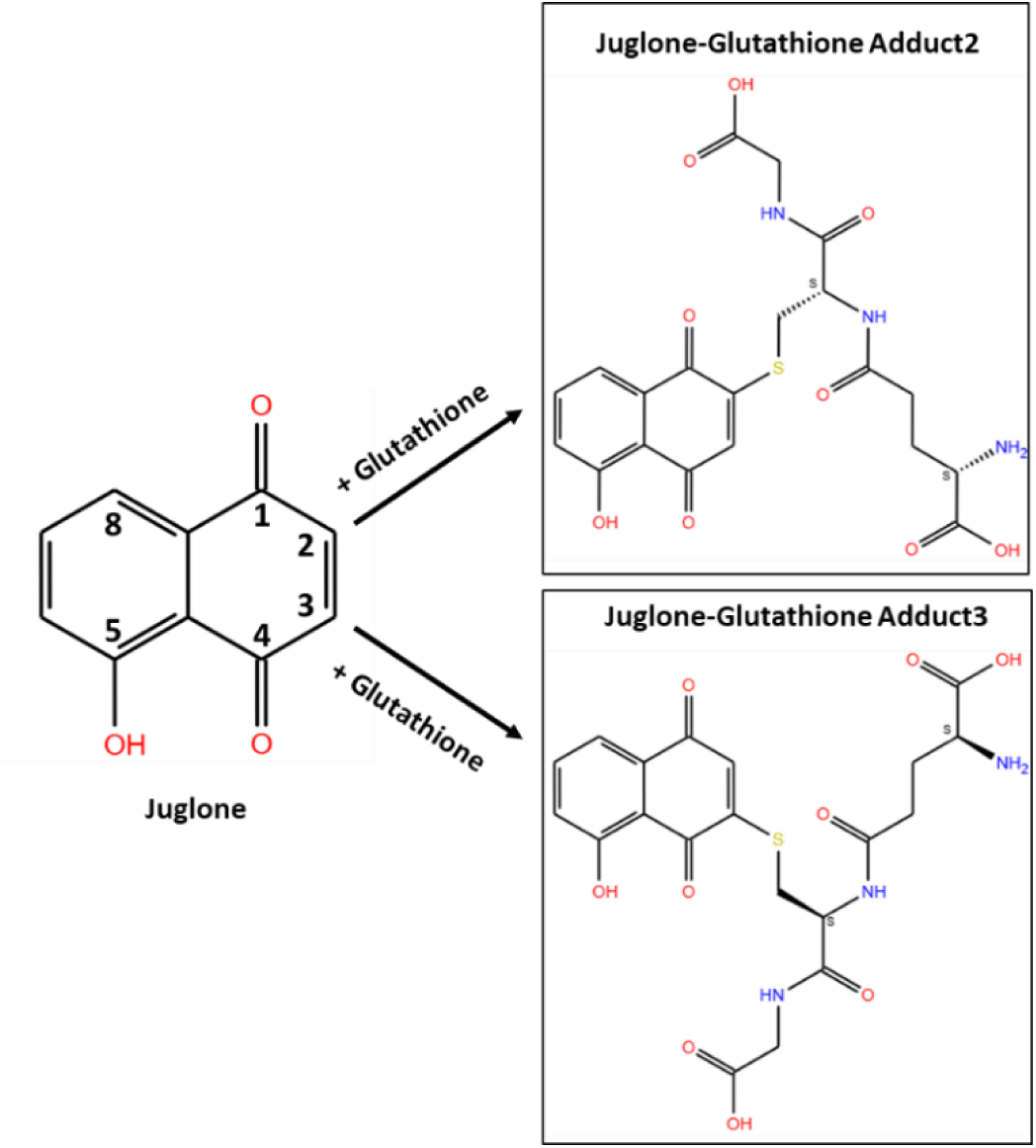
Schematic showing the formation of juglone-glutathione adducts drawn using ChemSketch program.

### Docking Methodology

The ChaC1-glutathione complex was used as the receptor for molecular docking where the binding site or the receptor grid center was determined from the coordinates of bound glutathione. The structure of juglone was downloaded from PubChem in SDF format and the adducts were modelled as per Scheme 1. Juglone and the adducts were then prepared for docking using LigPrep (15). Their ionization states were generated at pH 7.0 (±2.0) using Epik ionizer, and the lowest energy conformers were retained for each compound. The prepared compound conformations were subjected to Glide XP docking as described in the virtual screening section. The docking poses with the best scores were used to obtain the model systems followed by MD simulations.

### Molecular Dynamics (MD) simulations

The four model systems (ChaC1-Juglone, ChaC1-Glutathione-Juglone, ChaC1-Juglone-C2-Glutathione-adduct, and ChaC1-Juglone-C3-Glutathione-adduct) were subjected to protein preparation wizard of Maestro available with DESMOND MD simulation package (release 2018) (19) of Schrodinger. Structures were pre-processed to optimize the H-bond assignments at pH 7.00 using PROPKA. After pre-processing the model systems were solvated in a cubical water box (TIP3P water model) keeping 10 Å buffer space in x, y, and z dimensions. Neutralization of each system was done by adding appropriate counter ions while Na^+^ and Cl^-^ ions were added to maintain an ionic concentration of 0.15M. Under OPLS_2005 force field, each system was minimized with 5000 steepest descent steps followed by gradual heating from 0 to 300 K, under the NVT ensemble. The systems were thermally relaxed before the production run using the Nose-Hoover Chain thermostat method for 5 ns and 5 ns of pressure relaxation with the Martyna-Tobias-Klein barostat method. Final MD production run was performed for each system for 150 ns in NPT ensemble and OPLS_2005 force field. For the first model system (ChaC1-a-Glutathione) we already had MD trajectory for 100ns and we extended the simulations up to 150 ns. All trajectories were analysed with simulation event analysis and simulation interaction diagrams of DESMOND. Comparative analyses of the trajectories were performed to study the relative stabilities of the ligands bound to ChaC1. MMGBSA binding energies of the ligands were calculated from ten snapshots collected at equal intervals from the respective trajectories.

## Results

### Virtual Screening of two large Natural Products libraries for ChaC1 inhibitors and *in vitro* evaluation of the putative hits

In the absence of any known inhibitors of ChaC1, we decided to carry out a completely random *in silico* screen against the modelled ChaC1 structure of two large natural product libraries. These libraries included compounds of diverse molecular structures having drug-like properties in biologically relevant representations. Approximately 530,000 compounds from the Asinex library and 130,000 compounds from the ZINC natural products library were used. The virtual screening was performed as depicted (Figure 1) and described in the methods. The screen yielded a total of 102 hits.

**Figure 1.**
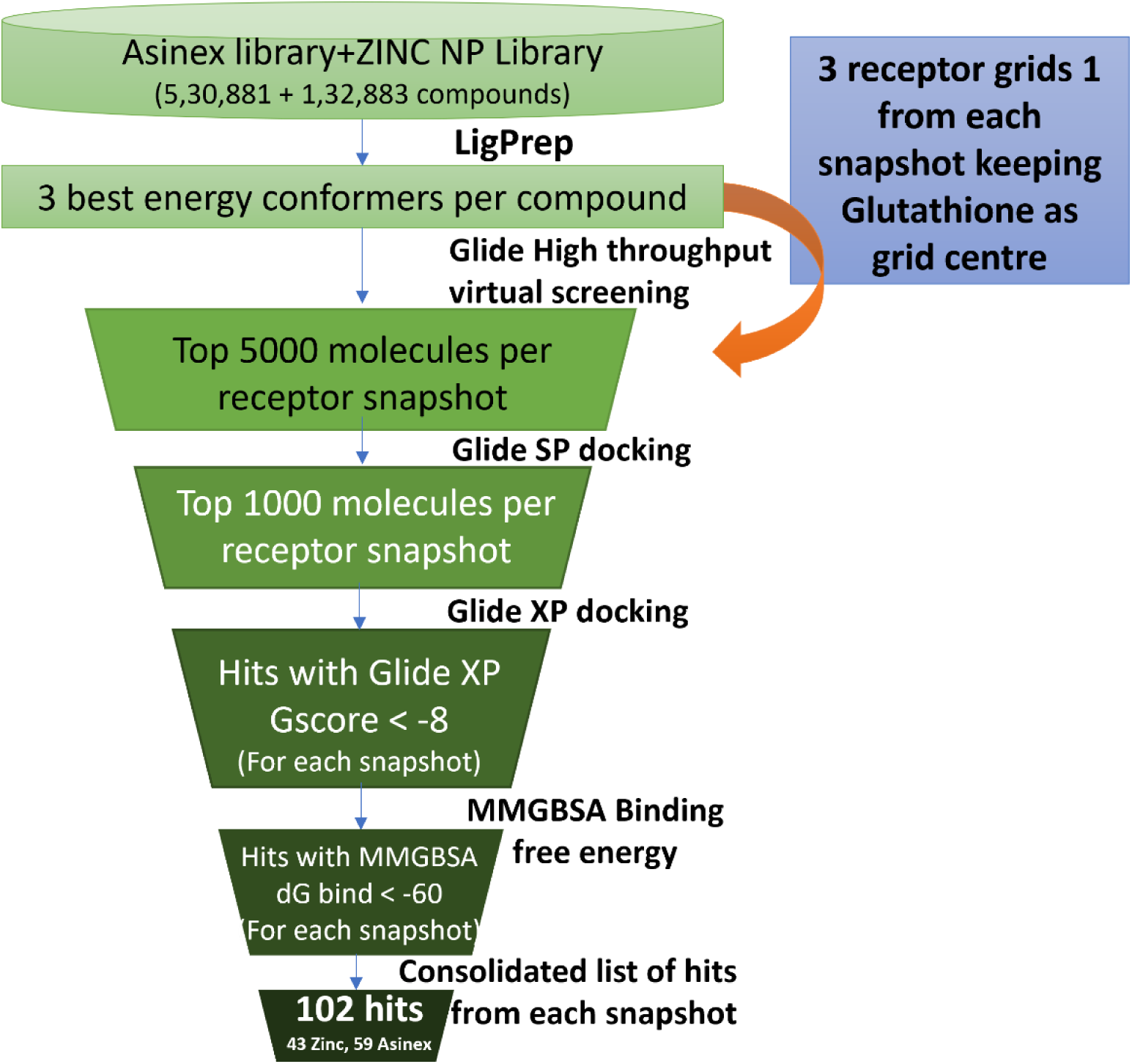
Virtual Screening workflow using Glide.

The top 10 screened hits formed complexes with ChaC1, with their binding free energies ranging between -96 to -80 kcal/mol, while their XP docking scores ranged between -8.4 to -8.1. Previous studies had demonstrated the essentiality of Y38, G39, S40, L41, D68, R72, E115 and Y143 residues for glutathione binding (14). All the screened hits also showed interactions with these essential glutathione-binding residues. The polar groups of these molecules also formed hydrogen bonds (H-bonds) with residues F36, Y38, G39, S40, L41, D68, R72, T83, E115, A116 and R181. The aromatic rings of these molecules were also found to make π-π stacking interactions with aromatic binding pocket residues like Y38, W43, Y143, and Y178. The aromatic rings of some of the hits also formed cation-π interactions with positively charged groups of R144 and R181 (Figure S1).

The top ten molecules were chosen for experimental testing based on their binding energies and interaction with key binding site residues. Nine out of these ten compounds were custom-made by a commercial source, MolPort (www.molport.com) (Figure 2A). These compounds at 100 µM were screened for their inhibitory activity against the human ChaC1 enzyme using the ChaC1p-Dug1p coupled assay. However, even at 100µM none of the compounds exhibited significant inhibition against the ChaC1 enzyme (Figure 2 B).

**Figure 2.**
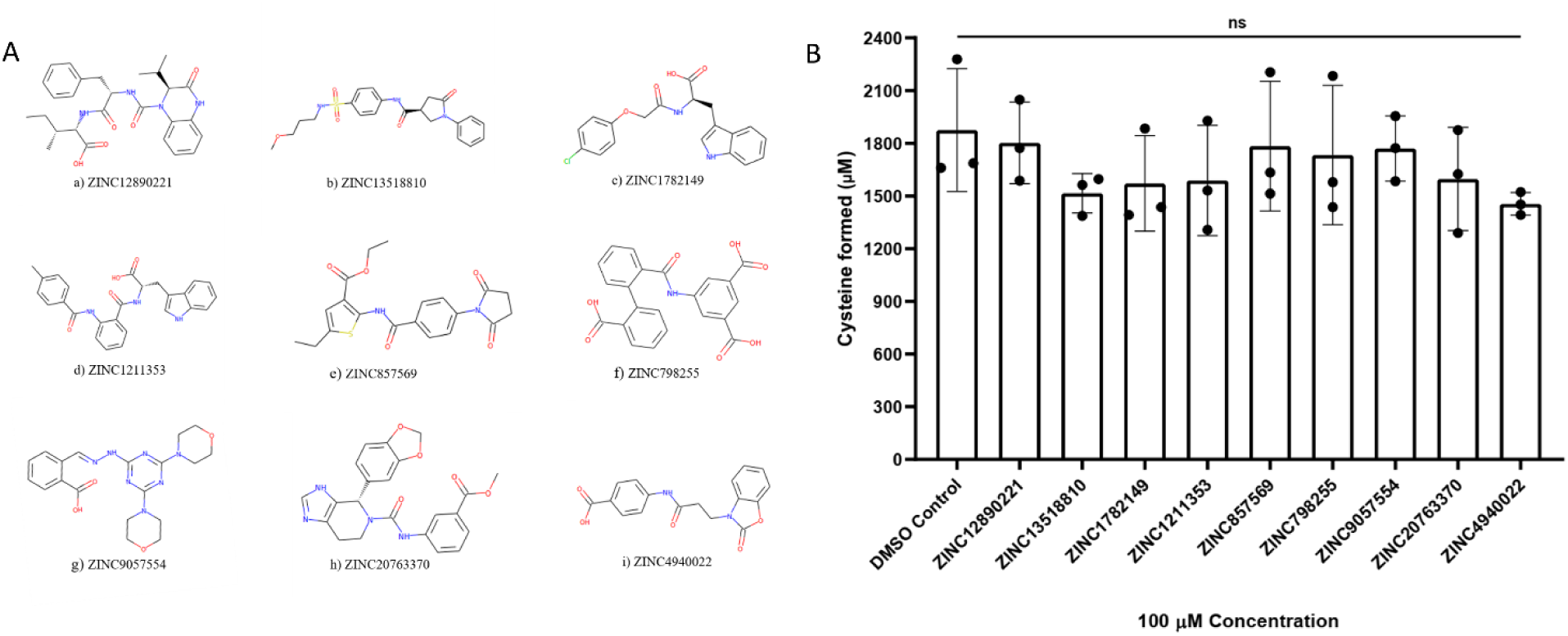
A Structure of 9 of the top 10 hits from the virtual screen custom synthesized for experimental evaluation. **a)** ZINC12890221-(2S,3R)-2-((S)-2-((S)-2-isopropyl-3-oxo-1,2,3,4 tetrahydroquinoxaline-1-carboxamido)-3-phenylpropanamido)-3-methylpentanoate **b)** ZINC13518810-(S)-N-(4-(N-(3-methoxypropyl)sulfamoyl)phenyl)-5-oxo-1-phenylpyrrolidine-3-carboxamide **c)** ZINC1782149-(R)-2-(2-(4-chlorophenoxy)acetamido)-3-(1H-indol-3-yl)propanoic acid **d)** ZINC1211353-(S)-3-(1H-indol-3-yl)-2-(2-(4-methylbenzamido)benzamido)propanoic acid **e)** ZINC857569-5-(2’-carboxy-[1,1’-biphenyl]-2-ylcarboxamido)isophthalic acid **f)** ZINC798255-ethyl 2-(4-(2,5-dioxopyrrolidin-1-yl)benzamido)-5-ethylthiophene-3-carboxylate **g)** ZINC9057554-(E)-2-((2-(4,6-dimorpholino-1,3,5-triazin-2-yl)hydrazono)methyl)benzoic acid **h)** ZINC20763370-(S)-methyl 3-(4-(benzo[d][1,3]dioxol-5-yl)-4,5,6,7-tetrahydro-3H-imidazo[4,5-c]pyridine-5-carboxamido)benzoate **i)** ZINC4940022-4-(3-(2-oxobenzo[d]oxazol-3(2H)-yl)propanamido)benzoic acid **B *In vitro* enzymatic assay for the virtual screening hits** 9 compounds procured from MolPort were dissolved in DMSO and their inhibition against the ChaC1 protein was evaluated at 100 μM concentration with 2 mM of substrate glutathione. The ChaC1p-Dug1p coupled assay was used to estimate the cysteine released as described in the methods. The experiment was done multiple times, with three technical replicates for each sample. The graph here corresponds to the representative data set plotted using the average of the three technical replicates and ± S.D. values. The p-value was determined using one-way ANOVA with multiple comparisons. ns: non-significant

### Glutathione analogues fail to function as efficient inhibitors of the human ChaC1 enzyme

Since active site inhibitor compounds often resemble the structure of the natural substrate, we decided to explore the possibility of γ-glutamyl tri peptides blocking the active site and acting as potential inhibitors for the human ChaC1 enzyme. Six glutathione analogs; γ-glu-leu-gly, γ-glu-met-gly, β-asp-cys-gly, β-asp-ser-gly, γ-glu-gly-gly, and γ-glu-ser-gly were custom synthesized (GenScript) for evaluation.

The glutathione analogues were also screened at a concentration of 100 µM each in the *in vitro* assay, but surprisingly all of them proved to be ineffective as inhibitors against the ChaC1 enzyme (Figure S2).

We also evaluated the inhibitory capability of N-glutaryl-L-alanine (GA) which is a deaminated analogue of its natural substrate, γ-glutamyl-L-alanine (20). This compound, GA, was identified as a potent inhibitor of the enzyme GGCT (γ-glutamyl cyclotransferase) which also belongs to the superfamily of cyclotransferases as ChaC1, and it also acts on γ-glutamyl amino acids. γ-GCT catalyzes the hydrolysis of γ-glutamyl-amino acids into 5-oxoproline and their corresponding amino acid (Figure S3) (21).

Since only the diester-modified form of GA (pro-GA, cell-permeable) was commercially available, we tested it against ChaC1 using the *in vitro* enzymatic assay. However, this pro-form appeared to have no significant effect on ChaC1 activity *in vitro* (Figure S3).

### Creation of two ‘opposite-acting’ yeast-based assays for high throughput screening for ChaC1 inhibitors

Since neither the *in silico* strategy nor the use of synthetic analogues could provide a tentative lead as an inhibitor against the ChaC1 enzyme, a random chemical compound library screening strategy was adopted. However, for this, we needed to develop a robust assay for the screening

Glutathione is an essential metabolite in yeast cells due to its role in Iron-Sulfur (Fe-S) loading in the mitochondria (22). Thus, in cells knocked out for glutathione synthesis in yeast, external glutathione needs to be added (23). In such cells (*gsh1Δ*), expression of ChaC1 can deplete the glutathione obtained from the medium and prevent the growth of cells.

Glutathione can also be a source of sulfur for yeast cells. The glutathione taken up can be degraded and the cysteine released can be used as the source of sulfur. Thus, in yeast cells that are organic sulfur auxotrophs (*met15Δ*), if the degradation of glutathione can be made dependent on ChaC1, then the functional expression of ChaC1 would allow the growth of cells that are dependent on glutathione as a sulfur source.

In the development of screens for ChaC1 inhibition, we have used these two properties of glutathione. The first screen thus employed a yeast strain that is a glutathione-deficient yeast where the first enzyme involved in glutathione biosynthesis, γ-glutamyl-cysteine synthetase (GSH1), is disrupted. As glutathione is essential for growth, yeast cells that cannot synthesize it, need to be provided external glutathione to allow growth. Even a small amount of glutathione is adequate for growth (1-10 μM) (23). When glutathione is added at such low concentrations, ChaC1 expression in these cells inhibits growth as it degrades the minimal amount of glutathione provided to them (1). Hence, the presence of a potential inhibitor of this enzyme would allow the growth of such glutathione auxotrophs (Figure 3A).

**Figure 3.**
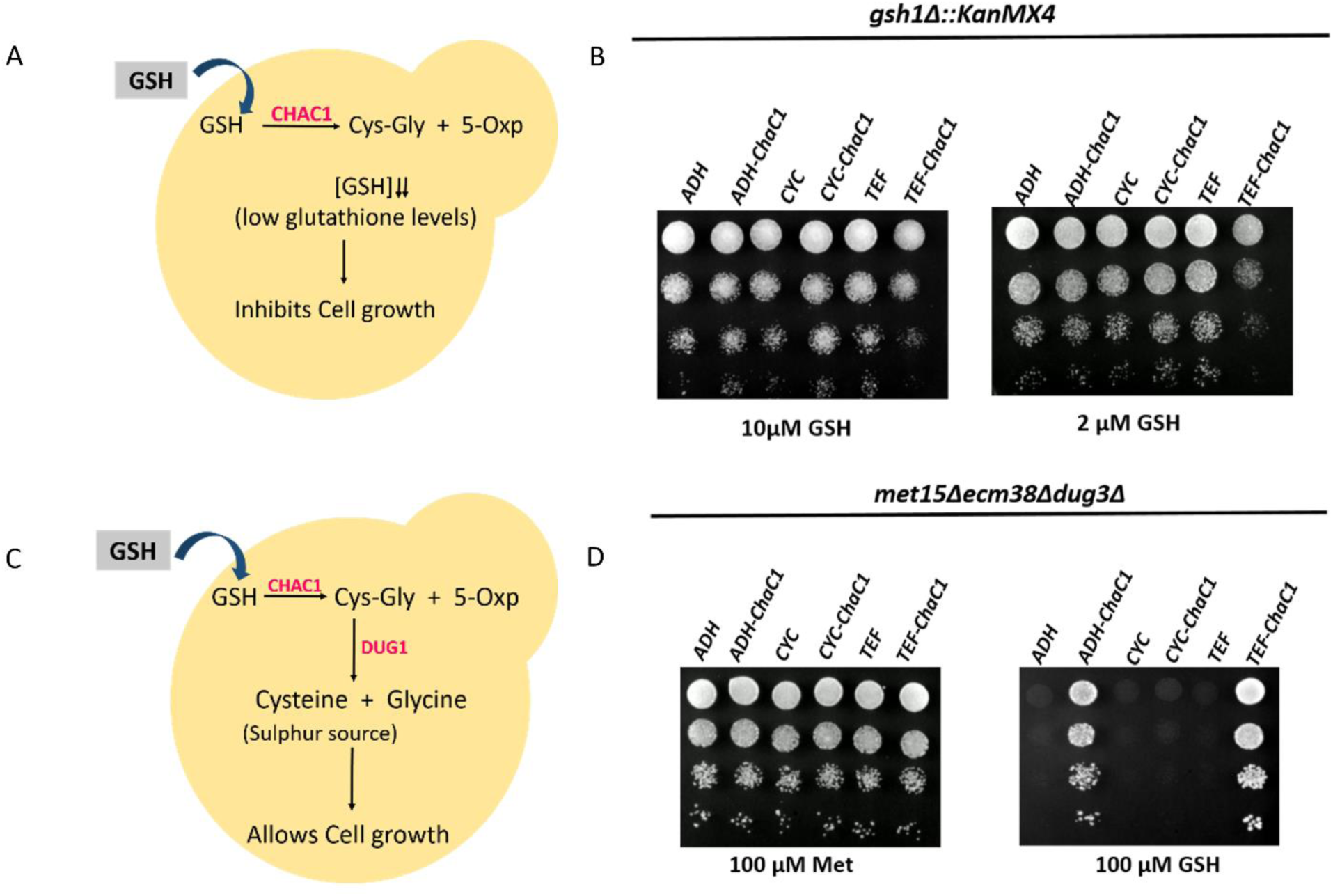
Optimization of human ChaC1 expression using promoters of varying strengths. Action of ChaC1 expression in the **A)** glutathione-deficient yeast, and **C)** sulphur auxotrophic yeast. **B)** and **D)** *S. cerevisiae* strains ABC1723 and ABC1976 were transformed with p416TEF-ChaC1, p416ADH-ChaC1, p416CYC-ChaC1, and their corresponding vector controls. The transformants were grown to exponential phase in a minimal medium, harvested, washed, resuspended in water, and serially diluted to give 0.1, 0.01, 0.001, and 0.0001 OD_600 nm_ of cells. Ten microliters of these dilutions were spotted on SD medium plates containing 200 μM of methionine or glutathione as a sulfur source. The photographs were taken after 48 hours of incubation at 30℃. The experiment was repeated twice, and a representative data set is shown.

The second assay was based on a yeast strain that was a sulfur auxotroph due to a deletion in the MET15 gene. This strain also carries deletions in the ECM38 and DUG3 genes. ECM38 encodes the glutathione-degrading enzyme γ-glutamyl transpeptidase while the DUG3 product is part of the Dug2p/Dug3p complex involved in cytoplasmic glutathione degradation in yeasts and fungi (24). Deletion of these genes makes these cells deficient in glutathione degradation. When ChaC1 is expressed in these cells, it degrades the glutathione which is provided externally, thereby providing cysteine as a sulfur source for their growth. In this case, an inhibitor of ChaC1 would suppress the growth of these cells (Figure 3C).

ChaC1 is a catalytically efficient enzyme. To evaluate the most appropriate expression level that could be used for the yeast-based screening assay, ChaC1 was cloned under yeast promoters of varying strengths in a single-copy vector. These included the strong TEF1 promoter, the moderately strong ADH1 promoter, and a weak CYC1 promoter. ChaC1 expressed downstream of these promoters was transformed into the glutathione deficient and the sulfur auxotrophic strains and then analyzed using the serial dilution growth assay on plates containing different concentrations of GSH.

ChaC1 expressed under the TEF1 promoter showed the best phenotypes in both screens (Figure 3B and 3D), so TEF1-ChaC1 was chosen for both screens.

Both these strains were further engineered to be drug-sensitive by creating a deletion of the Pdr5 efflux pump (as described in methods). The deletion was validated using the cycloheximide sensitivity assay (Figure S4).

### High throughput screening for ChaC1 inhibitors: optimization, automation, and hits identification

The two screens were initially established on solid agar plates. However, we needed to demonstrate the screens in a 96-well plate format to enable cost-effectiveness and increased throughput. We accordingly established these assays in low-volume (200 µl) liquid cell cultures that were optimized for glutathione concentration, incubation time, shaking, and measurement intervals in a plate reader and these are described in the methods. We also determined that 2.5% DMSO concentrations (for compound solubilization) did not hamper the screens (Figure S5). To determine the robustness of the screen the statistical parameter, Z-prime (Z’) was determined (25) and at 40 hours of growth the values obtained were very significant (for Screen I =0.95 and for Screen II = 0.91) which suggested that the screens we developed were robust and would be compliant for a high throughput setting (Figure S6).

After the successful manual optimization of the assays in a plate reader, automated compound library screening was performed against a Spectrum library using a Tecan robotic liquid handling system. The Spectrum library was chosen because this library contains a total of 2320 compounds of diverse shapes and structures, and is based on commonly occurring compounds with established biological functions. In the screen against the glutathione-deficient yeast strain (Screen I), where any compound inhibiting ChaC1 would permit the growth of these cells (greater than three times the standard deviation of the mean library effect), were considered for further evaluation (Table S2). However, as several compounds showed this effect, we prioritized the ones displaying higher growth which could be marked distinctively. A total of 14 hits were initially obtained based on the OD_600 nm_ values obtained at the end of 36 hours of the growth experiment (Figure 4).

**Figure 4.**
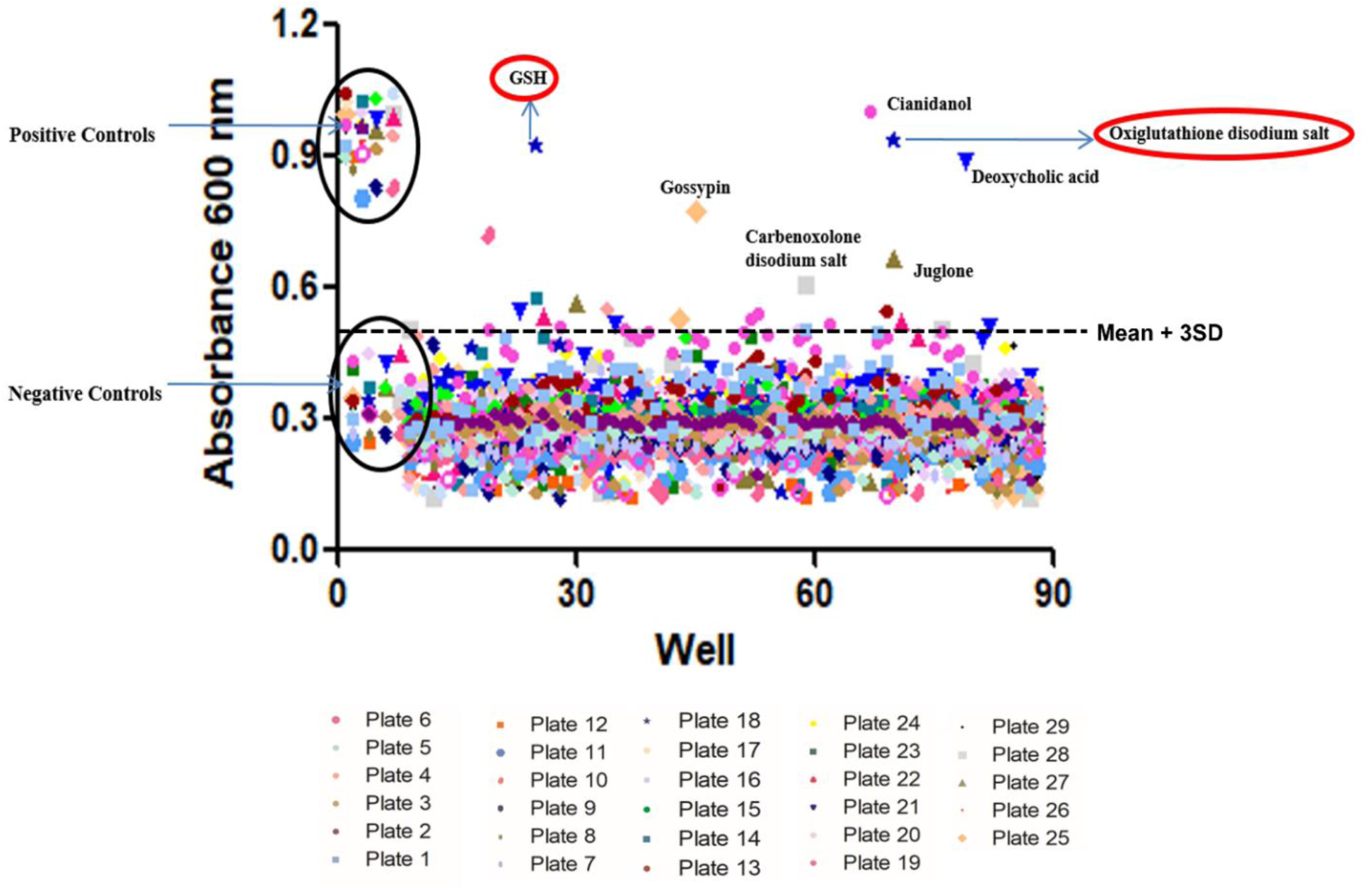
High-throughput screening of Spectrum library using Screen I (*gsh1Δ* yeast strain) *S. cerevisiae* strain ABC 6303 transformed with ChaC1 and the corresponding vector control was grown according to the growth conditions optimized for High-throughput screening assay (Methods). 200 μL (0.15 OD_600 nm_) of ChaC1 expressing yeast cell culture was dispensed in six 96-well microtiter plates and 100 μM (10 μL) of each compound from the Spectrum library was added using a liquid handling system. Controls with only DMSO were also added to each plate. The plates were incubated at 30℃ with continuous shaking in the incubator and the growth was monitored for 36 hours. Endpoint OD_600 nm_ were obtained for each compound and plotted using graph pad prism software. The dotted line indicates three standard deviations above the mean effect.

Growth curves for these apparent hits were generated which helped us identify and omit some coloured compounds that were false positives and did not affect the cell growth (data not shown). The compounds (and the OD_600 nm_ reached) of these positive hits included Cianidanol (OD_600 nm_ =1), Juglone (OD_600 nm_ =0.66), Deoxycholic acid (OD_600 nm_ =0.89), Carbenoxolone (OD_600 nm_ =0.58), Gossypin (OD_600 nm_ =0.77)(Figure S7). The hits also included Glutathione (OD_600 nm_ =0.93) and Oxidized glutathione (OD_600 nm_ =0.92) as they were also part of the Spectrum library, and were an internal validation of the screen (Table S2).

The screening procedure was repeated with the sulfur auxotrophic strain where an inhibitor of ChaC1 would impede the growth of yeast cells. The hits that allowed growth in the first screen should have shown opposite effects in Screen II. However, in the second screen, hundreds of compounds led to growth inhibition (Figure S8). As it appeared that the majority could be off-targets or have general cytotoxicity, we decided to focus on the hits from Screen I for further *in vitro* enzymatic validation.

### *In vitro* analysis of hits from HTS reveals juglone as a genuine and efficient inhibitor of ChaC1

The five compounds obtained as hits were procured afresh and evaluated using the ChaC1p *in vitro* enzymatic assay (24). These compounds were used at a concentration of 100 µM against ChaC1 with 2 mM substrate (GSH). Interestingly, only one compound, juglone, was able to completely inhibit ChaC1 activity at this concentration (Figure 5).

**Figure 5.**
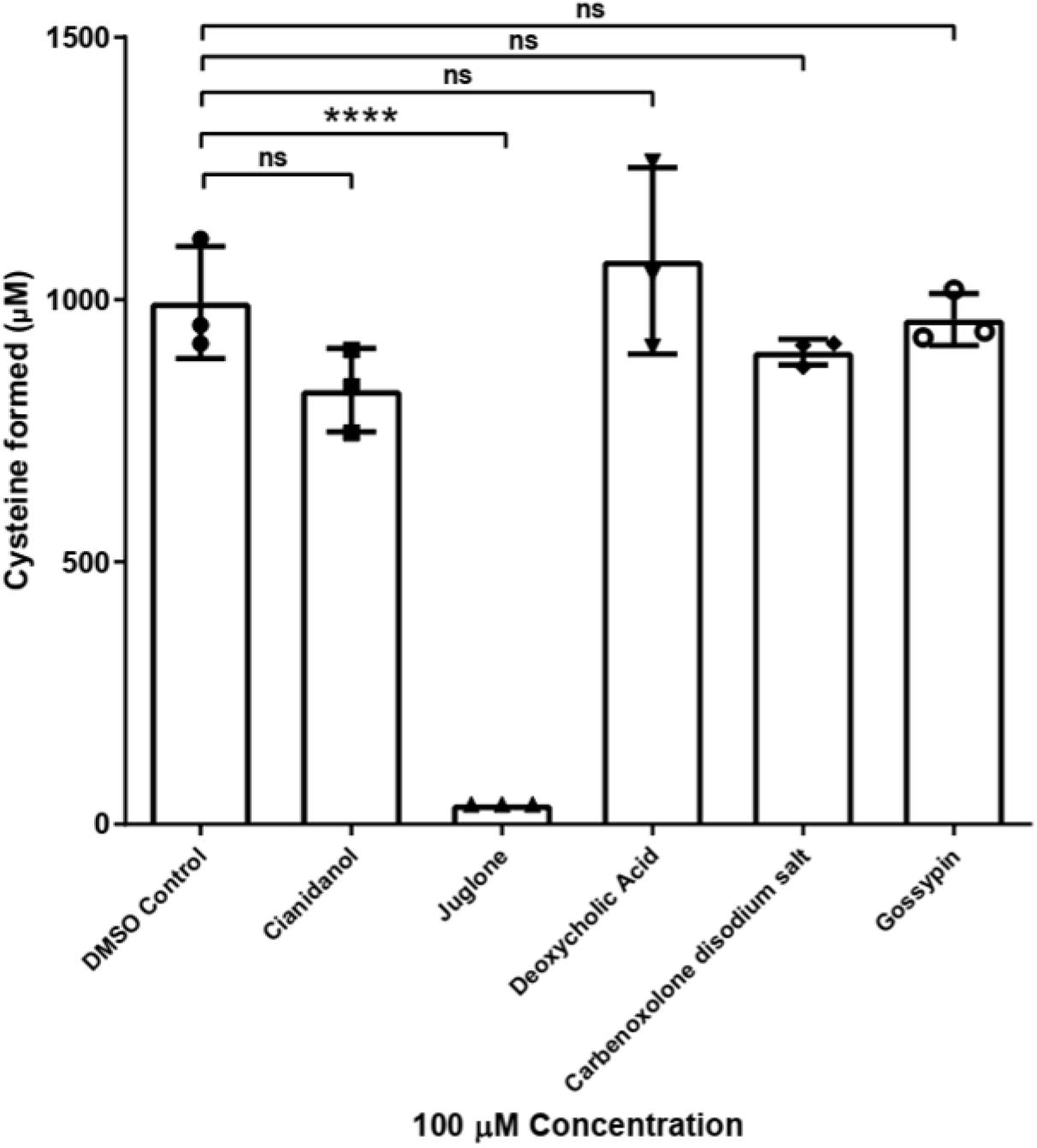
*In vitro* enzymatic assay of hits obtained from Screen I. Cianidanol, Juglone, Deoxycholic Acid, Carbenoxolone disodium salt, and Gossypin were evaluated for their inhibition against the ChaC1 protein at 100 μM concentration with 2 mM of substrate glutathione. The ChaC1p-Dug1p coupled assay was used to estimate the cysteine released as described in methods. The experiment was done thrice, along with three technical replicates for each sample. The graph here corresponds to the representative data set plotted using the average of the three technical replicates along with ± S.D. values. The p-value was determined using one-way ANOVA with multiple comparisons. ns: non-significant, p>0.05, *p<0.05, **p<0.01, ***p<0.001, ****p<0.0001

To estimate the inhibition profile of juglone, a range of concentrations between 0.25 µM to 100 µM were tested against 2 mM glutathione. Our results indicated that juglone had IC_50_ values of 8.7 µM when using 2 mM substrate concentrations (Figure 6). We also evaluated higher glutathione concentrations, since the cytosolic glutathione concentrations have been reported to go up to 10 mM in mammalian cells (26). When the inhibition was carried out with glutathione at 4 mM, juglone had an IC_50_ value of 21.1 µM respectively. Thus, even with the substrate at approximately 250 to 300-fold higher concentrations, juglone inhibition was significant (Figure S9 B). Interestingly, juglone at 100 μM concentration was also able to inhibit the ChaC1 activity by 50% when tested at even higher glutathione concentrations (10 mM glutathione) (Figure S9 C). This indicated that juglone was indeed a potent inhibitor of ChaC1.

**Figure 6.**
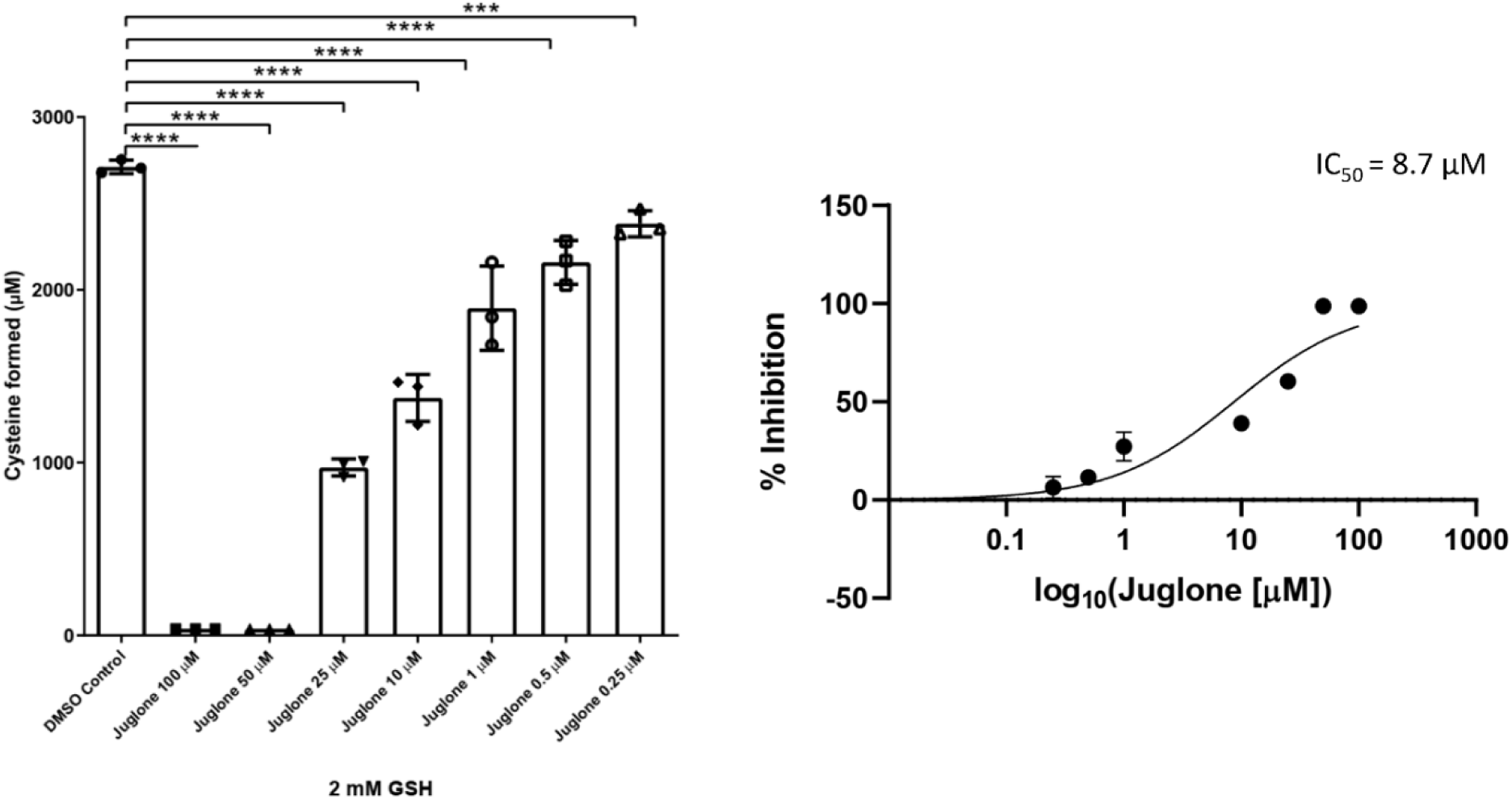
Determination of IC_50_ of juglone against ChaC1 enzyme. Different concentrations of juglone were evaluated for their inhibition against the ChaC1 protein. The ChaC1p-Dug1p coupled assay was used to estimate the cysteine released as described in methods. The IC_50_ value for juglone was determined against 2 mM GSH. The experiment was done thrice, along with three technical replicates for each sample. The graph here corresponds to the representative data set plotted using the average of the three technical replicates along with ± S.D. values. The p-value was determined using one-way ANOVA with multiple comparisons. ns: non-significant, p>0.05, *p<0.05, **p<0.01, ***p<0.001, ****p<0.0001

### Juglone is also an efficient inhibitor of the ChaC1 paralog, ChaC2

ChaC2 is a homolog of ChaC1, and higher eukaryotes including humans have both ChaC1 and ChaC2 present in their cytosol responsible for glutathione degradation. The human ChaC2 has a 50% identity with the human ChaC1 protein. To examine if ChaC2 is also inhibited by juglone, we evaluated the purified human ChaC2 protein activity *in vitro* in the presence of juglone. Our results revealed that juglone could also inhibit ChaC2 efficiently. Although ChaC1 was completely inhibited by juglone at 50 μM concentration, in the case of ChaC2, a slightly higher concentration (100 μM) was required (Figure 7).

**Figure 7.**
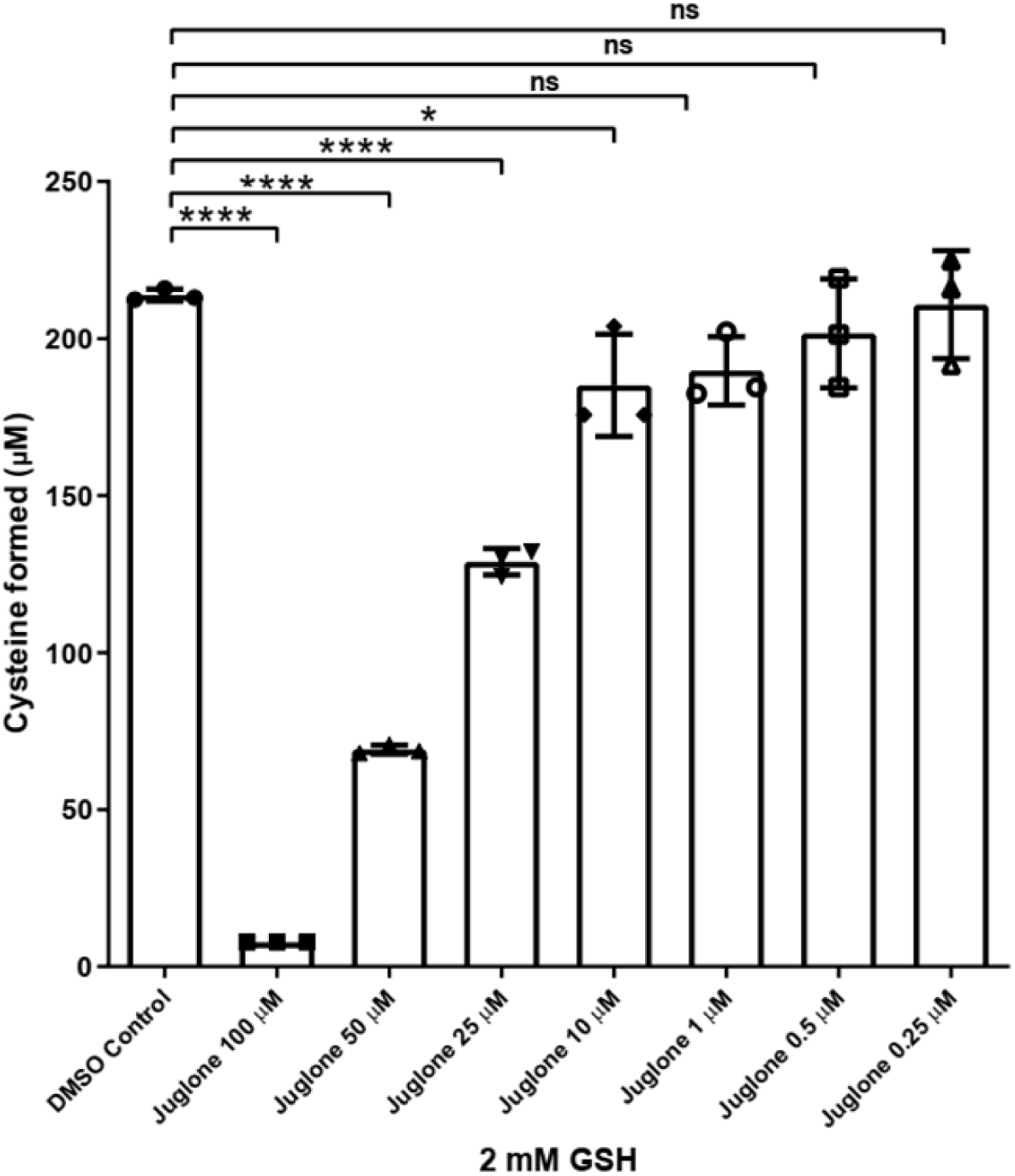
Inhibition ChaC2 by juglone. Different concentrations of juglone were evaluated for their inhibition against the ChaC2 protein with 2 mM substrate glutathione. The ChaCp-Dug1p coupled assay was used to estimate the cysteine released as described in methods. The experiment was done twice, along with three technical replicates for each sample. The graph here corresponds to the representative data set plotted using the average of the three technical replicates along with ± S.D. values. The p-value was determined using one-way ANOVA with multiple comparisons. ns: non-significant, p>0.05, *p<0.05, **p<0.01, ***p<0.001, ****p<0.0001

As the ChaC1p and ChaC2p enzymatic assays were coupled assays involving the cys-gly dipeptidase enzyme Dug1p (coupled to ChaC1p or ChaC2p), it was possible that the compound juglone could be inhibiting this second enzyme and not ChaC1 or ChaC2. To verify the inhibitory effect, if any, of juglone on Dug1p, a Dug1p activity assay was performed using cys-gly as a substrate. Our results indicated that juglone had no inhibitory effect against Dug1p (Figure S10).

### Juglone does not inhibit ChaC1 by conjugation to its cysteines, as seen with the strong inhibition of a cysteine-free ChaC1 mutant by juglone

Juglone is a naturally occurring naphthoquinone found in walnuts and has long been in usage. The mechanisms by which juglone has been shown to act within cells appears to involve inhibition through the formation of adducts either with sulfhydryl proteins, with cellular GSH or through the continuous formation of semiquinone radicals through the action of NADH/NADPH dependant reductases that generate deleterious ROS in the cell (27).

At the protein level, juglone has been shown to specifically inhibit a few purified enzymes by covalently modifying their active site cysteine residues. These include the Peptidyl-prolyl isomerase (Pin-1) in humans at the Cys113 residue (28), the Sortase A enzyme from *S.aureus* at the Cys183 residue (29), and the RNase P enzyme from *A. thaliana* at the C353 and C373 residues (30).

As the ChaC1 enzyme has GSH as its substrate (added at 2 mM), two possibilities of juglone inhibition of ChaC1 immediately suggest themselves. The first is depletion of the GSH substrate by conjugation to GSH. The second is the inactivation of ChaC1 by modification of cysteines. However, the formation of an adduct with GSH is unlikely to lead to sufficient depletion of GSH to be the cause of inhibition for the ChaC1 enzyme. This is because the reaction mixture contains nearly 200 times higher amounts of the substrate (GSH) than the inhibitor (juglone at 10 μM). Under these conditions, although adduct formation is quite possible, a substantial amount of substrate remains available for the ChaC1 enzyme to catalyze the reaction.

The inhibition of ChaC1 by covalent modification of the cysteine residues of the ChaC1 protein, therefore, presented itself as a likely possibility.

The human ChaC1 enzyme contains 5 cysteine residues (Figure S12 A). Although none of these cysteines is in the active site, the probability of the modification of one of the cysteines by juglone leading to its inactivation, as seen in the case of Pin-1, Sortase, and RNase P enzymes, seemed a probable mechanism of inhibition. To test this hypothesis, we created a cysteine-free ChaC1 mutant and analyzed the effect of juglone on its activity. The cysteine-free protein is known to retain up to 90% activity (Figure S11). Our results demonstrated that juglone was able to strongly inhibit the activity of the cysteine-free mutant. The inhibition against the cysteine-free enzyme seemed as effective as that seen against the WT protein (Figure S12 B). This indicated that the mechanism of inhibition of juglone on ChaC1 was independent of cysteines, unlike in the case of Pin-1, Sortase, and RNase P enzymes.

### Juglone is not an active site competitive inhibitor of the ChaC1 enzyme

The inhibition of ChaC1 by juglone was not linked to its cysteine residues, nor did it appear to act through GSH depletion since GSH in the reaction was in far excess. At the protein levels, these are the two known mechanisms of juglone action that did not seem to be operating in the present case. As juglone was also inhibiting both ChaC1 and ChaC2 efficiently, it appeared that juglone might be inhibiting ChaC1 at its active site through competitive inhibition with glutathione. To evaluate this, we carried out competition kinetics of juglone and glutathione with ChaC1. As can be seen from the figure (Figure S13), we did not observe competitive kinetics but instead observed mixed kinetics, which indicated that juglone inhibition of ChaC1 was not competitive with glutathione as was initially expected.

To further confirm this observation, we carried out MD simulations. A Juglone-ChaC1 model was subjected to 150 ns MD simulations followed by analysis of the trajectories (Figure S14) as described in the methods.

The ChaC1-Juglone complex was designed to examine if juglone binds directly to the glutathione binding site as a competitive inhibitor. As per initial docking results, juglone seemed to bind exactly in the glutathione binding region making H-bond interactions with Y38 and Y121. However, within 1ns of MD simulation, it shifted to an adjacent site, remained there up to 140 ns, and after 140 ns completely dissociated from ChaC1 as evident from the sudden rise of RMSD of juglone (red line) with respect to the protein up to 18 Å as seen in figure 9 A. This indicated juglone might not be competing with glutathione for the binding site, in agreement with the kinetic studies (Video S1).

To find out whether juglone has an affinity for any other binding site other than the glutathione binding site, we generated a slightly different model system, where glutathione was bound to ChaC1 protein at its binding site, and juglone was blindly docked to this system. We observed that juglone was not able to stably bind to any of the nearby pockets of ChaC1. The RMSD of juglone with respect to the ChaC1 protein was found to be as high as 60 Å (Figure 9 A grey dotted line). By the end of the simulations, juglone completely dissociated from ChaC1. It was interesting to note that, juglone binding did not affect the Glutathione-ChaC1 interactions during the simulations (Figure 9 A grey solid line). Thus, allosteric inhibition by juglone also seemed unlikely from these observations (Video S2).

### Molecular Dynamic simulations suggest that juglone-glutathione conjugates could be strong binders to ChaC1

In the absence of direct inhibition of juglone with ChaC1 either by competitive or allosteric mechanisms, the other possibility that remained to be examined was whether the juglone-glutathione conjugate adduct that can form during the interaction of juglone with glutathione can act as a competitive inhibitor of ChaC1. Many previous studies have reported evidence of the formation of such glutathionyl quinone or naphthoquinones adducts (31,32) .The C-2 and C-3 in juglone, can react with nucleophiles like glutathione to form adducts. We constructed two such adducts (Juglone-C2-Glutathione-adduct and Juglone-C3-Glutathione-adduct) *in silico* as shown in Scheme1 and docked them to ChaC1 at the glutathione binding site to generate the last two model systems. From both the docked complexes, it is observed that the glutathione parts of the adducts make H-bond and/or electrostatic interactions with the key glutathione binding site residues S40, R44, R72, R114, E115, and S172. The juglone part of the adduct is surrounded by hydrophobic residues like Y110, Y113, and F171 in both the complexes while an additional π-π stacking with Y110 was observed in the C3-adduct complex. MD trajectories of both the adducts complexed with ChaC1 also showed very stable interaction profiles (Figure 8) indicating stable binding in the ChaC1 binding site. However, the C3-adduct appeared to be more stable as the oxygen atoms of the C4 carbonyl group and C5-hydroxyl groups made additional water bridges with R44 and D46 respectively.

**Figure 8.**
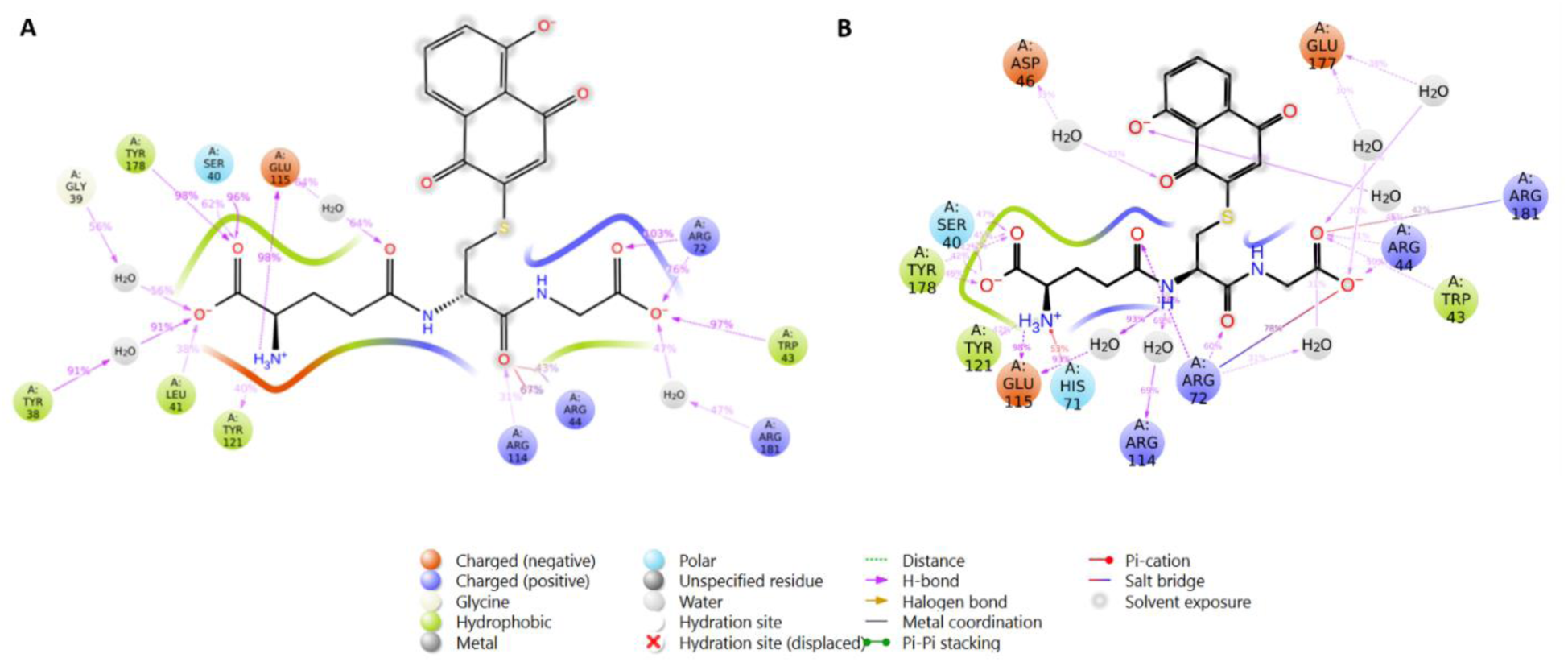
Interaction profiles of A) Juglone-C2-Glutathione-adduct and B) Juglone-C3-Glutathione-adduct during 150 ns MD simulations.

Looking at the stabilities of these adduct complexes we performed further comparison of their docking scores (before MD simulations) and the average MMGBSA binding energy throughout the MD simulations. The average MMGBSA binding energy was observed to be lower (better) in case of the Juglone-C3-Glutathione adduct (-55.07 kcal/mol) (Figure 9 B) than that of Juglone-C2-Glutathione adduct (-47.76 kcal/mol) (Video S3). This observation shows that the C3 adduct is a better binder than the C2 adduct. Therefore, our molecular dynamics study supports the hypothesis that juglone can form adducts with glutathione at positions C-2 and C3. Of these two adducts, the adduct with glutathione at the C3 position seemed to be a more effective inhibitor of ChaC1 (Video S4).

**Figure 9.**
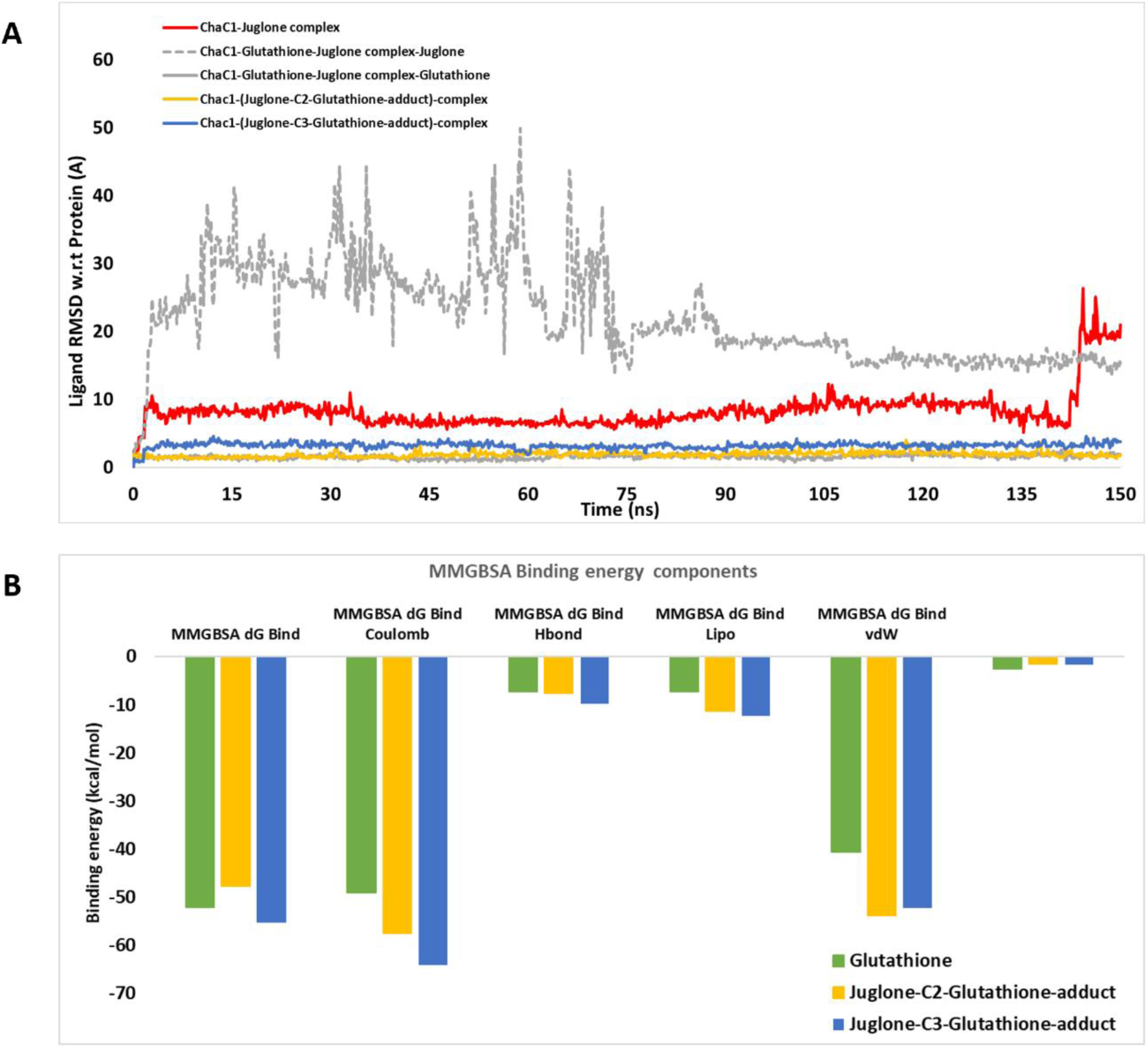
A) RMSD of the ligands with respect to ChaC1, B) Average MMGBSA binding energy and its components in different complexes during 150 ns MD simulations.

Juglone-glutathione conjugates were not available commercially, and we were also unable to custom synthesise them to confirm this possibility experimentally. In the absence of these specific conjugates, we sought to examine whether ChaC1 could be inhibited by other glutathione conjugates. We experimentally evaluated two commercially available glutathione conjugates, S-methyl glutathione, and glutathionyl glutathione (oxidized glutathione). However, neither of them, was able to inhibit ChaC1 to any significant extent (data not shown).

### Evaluation of other naphthoquinones and juglone analogues against ChaC1 reveals plumbagin as a second inhibitor against ChaC1

The MD simulations suggested that the Juglone-GS conjugate formation was responsible for the inhibition of ChaC1 (and not juglone *per se*) since other known mechanisms were eliminated. We therefore decided to examine how other analogues of juglone might behave since the reactivity of these naphthoquinones towards glutathione as well as the ability of the formed conjugate adducts to bond with the amino acid residues of ChaC1 were both important for the activity against the ChaC1 enzyme.

Several analogues were evaluated for their inhibitory activity against ChaC1 (Figure 10).

**Figure 10.**
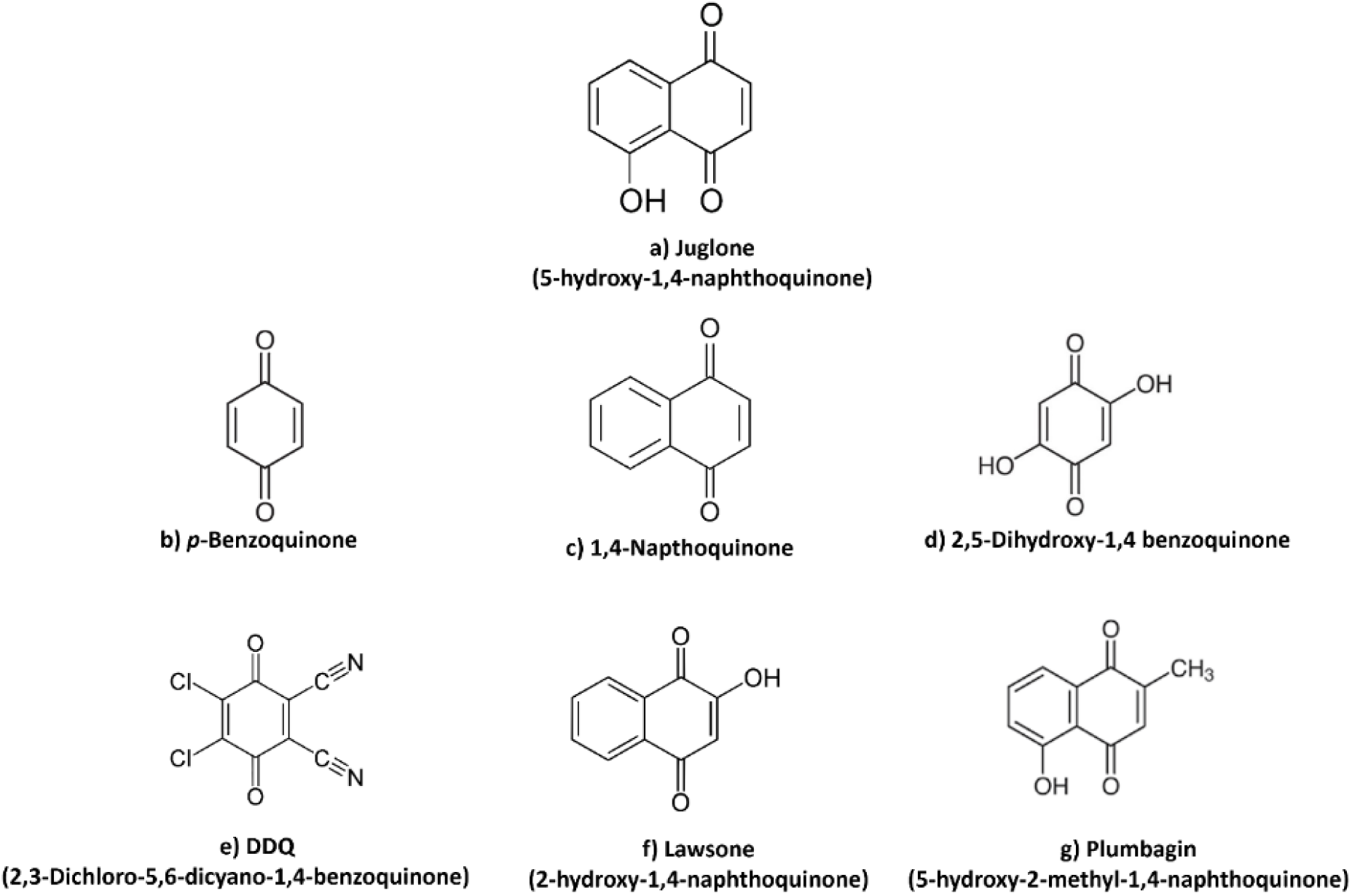
Chemical structures of related naphthoquinones and juglone derivatives.

Among those tested, the compound that showed virtually no inhibition was 2,5-dihydroxy 1,4 benzoquinone (Figure 11) and was the one predicted to be the least reactive. Among the other compounds tested, p-benzoquinone also showed lesser inhibition. Although this compound would be expected to be significantly reactive towards glutathione, it is also expected that the adduct formed would have a high propensity to revert back to the initial substrates by elimination through a retrograde Michael addition. This might be one of the factors that could explain the relatively lower inhibition of this compound.

**Figure 11.**
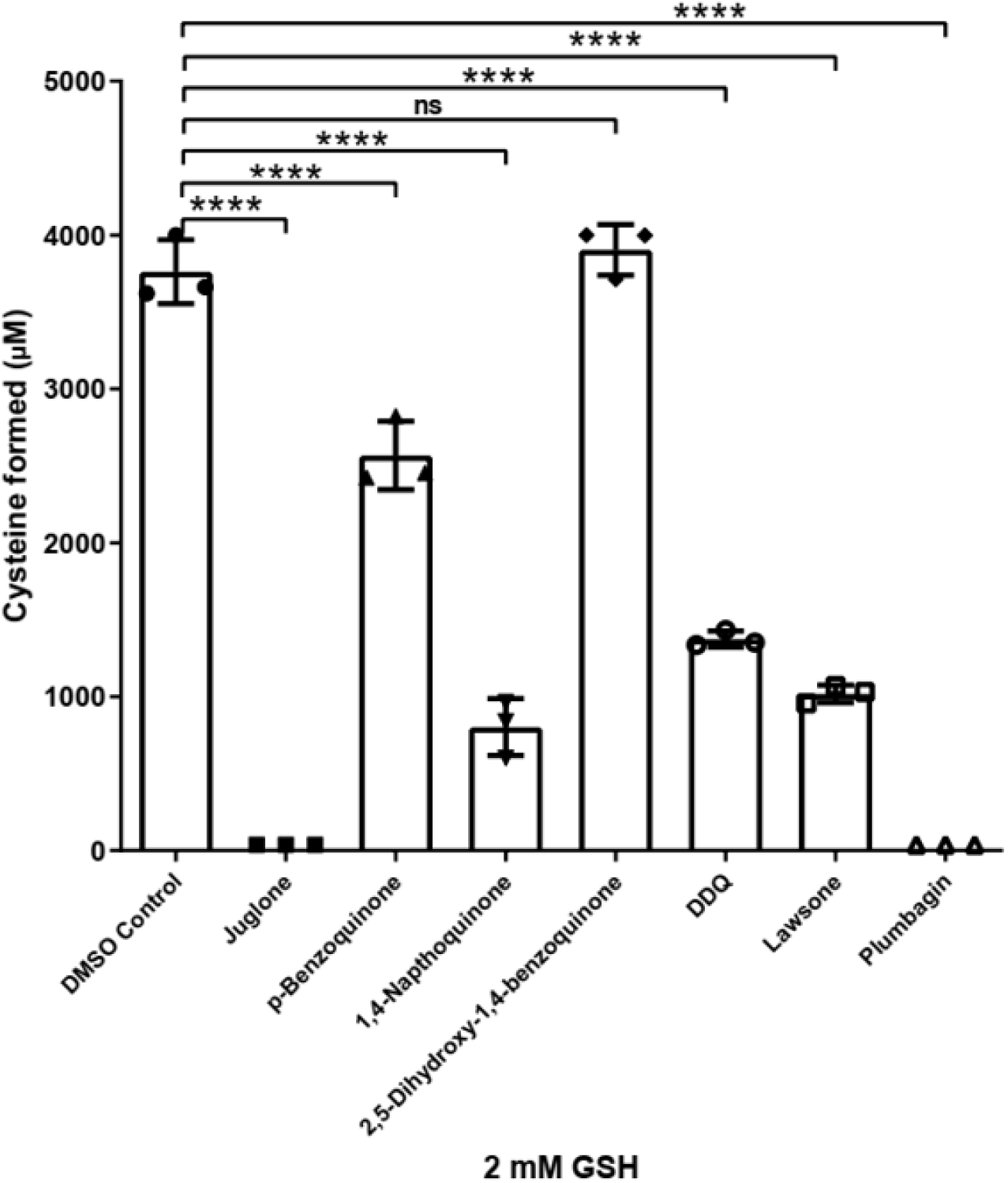
*In vitro* enzymatic assay of other naphthoquinones. p-benzoquinone,1,4-naphthoquinone, 2,5-dihydroxy-1,4 benzoquinone, DDQ (2,3-Dichloro-5,6-dicyano-1,4-benzoquinone), Lawsone, and Plumbagin were evaluated for their inhibition against the ChaC1 protein at 100 μM concentration with 2 mM of substrate glutathione. The ChaC1p-Dug1p coupled assay was used to estimate the cysteine released as described in methods. The experiment was done thrice, along with three technical replicates for each sample. The graph here corresponds to the representative data set plotted using the average of the three technical replicates along with ± S.D. values. The p-value was determined using one-way ANOVA with multiple comparisons. ns: non-significant, p>0.05, *p<0.05, **p<0.01, ***p<0.001, ****p<0.0001

In general, compounds that contained the quinone structure, and only a single aromatic ring such as p-benzoquinone, 2,5-dihydroxy 1,4 benzoquinone, and DDQ (2,3-Dichloro-5,6-dicyano-1,4-benzoquinone) showed very limited inhibition even at 100 μM (Figure 11).

Among those compounds that had two aromatic rings along with the quinone structure, 1,4 naphthoquinone and lawsone showed slightly more inhibition. However, even lawsone which had a hydroxyl group on the same quinone ring, did not show substantial inhibition. Only plumbagin (5-Hydroxy-2-methyl-1,4-naphthoquinone), which had two aromatic rings with the hydroxyl and quinone placed very similar to juglone showed almost identical inhibition to juglone.

Based on these investigations it appears that both the reactivity and the structure of the naphthoquinone structures were important. The reactivity would be important for the ability to form the glutathione conjugate, while other moieties and functional groups of the molecule would also be important for the ability to form strong binding interactions with the ChaC1 protein.

As plumbagin showed strong inhibition similar to juglone, we also examined how it would inhibit the cysteine-free ChaC1 molecule. Interestingly, here also we found that plumbagin could strongly inhibit the cysteine-free molecule, indicating that it was also functioning in a manner very similar to juglone (data not shown).

## Discussion

There is an important need to find inhibitors of ChaC1, a recently discovered enzyme that efficiently degrades glutathione in the mammalian cytosol. ChaC1 is highly induced under stress conditions. As it acts only on reduced glutathione, its induction depletes reduced glutathione pools leading to a very oxidizing environment. However, there are multiple challenges to identifying efficient inhibitors of ChaC1. These challenges include the lack of crystal structure of ChaC1, the absence of prior inhibitors, and most importantly the high Km of ChaC1 for glutathione. Considering these challenges, we have taken a very comprehensive approach towards inhibitor discovery.

The active site focussed approach that we initially adopted involved the virtual screening of large compound libraries. However, this was unable to provide us with any hits that could be confirmed when we tested them experimentally. The reasons for the failure of the virtual screen to yield even a tentative lead are not clear. One reason could be the static nature of the ChaC1 structures considered for the virtual screening. Though the structures were obtained at equal intervals from the MD trajectories of ChaC1-glutathione complexes, the binding of the hit molecules might not be precise. False positive hits at this stage could have led to errors in the hits obtained. Secondly, it is possible that more compounds need to be screened from amongst the hits to increase the chances of success. Finally, it is possible that these compounds could not compete with the 20-fold excess of glutathione substrate that is used in the enzymatic assays. The latter seems to be the most likely possibility since even custom-synthesized peptides that were substrate analogs (of GSH) failed to be effective as ChaC1 inhibitors.

Since we were always concerned that the high substrate concentrations (2 mM) might interfere with an active site inhibitor, we also carried out a chemical library screening approach. This approach involved the development of two novel yeast-based high throughput assays where heterologous expression of the human ChaC1 enzyme led to the growth of yeast cells in one assay and cessation of growth in the other assay. These would then be screened against chemical libraries. This approach would be active-site independent. This would be able to identify and capture inhibitors that could be either active site, non-active site inhibitors, or allosteric. Deploying these screens in a high throughput format and screening the widely used Spectrum library that includes biologically active compounds with significant structural diversity, we could finally identify one compound, juglone, that was able to completely inhibit the ChaC1 activity even at 50 μM.

Interestingly, juglone is a naphthoquinone, found in walnuts, and has been a part of old Chinese folk medicine for many years (33,34). It is also known to exhibit anti-tumor effects against various types of cancers (35,36). Juglone has been found to be cytotoxic in cell lines at even 20 μM concentrations. This might be a consequence of its ability to generate ROS through the formation of semiquinones in a process called redox cycling (31). However, as the ChaC1 enzyme is upregulated in many cancers (11), part of the cytotoxicity could be owing to its ability to inhibit ChaC1 that are important for the survival of these cell lines. The findings described here could thus shed more light on some of the earlier studies of juglone on cancer cells.

Since juglone was shown to inhibit a cysteine-free ChaC1, and since it could inhibit both purified ChaC1 and ChaC2 *in vitro*, it appeared that juglone might be acting in a mechanism not normally observed or previously reported for Juglone - through active site inhibition. However, neither the rigorous MD simulations carried out up to 150 nanoseconds, nor the competitive kinetic studies done with GSH supported this possibility.

This led us to the possibility that inhibition was occurring through adduct formation, followed by inhibitory action of the adduct. Juglone and other naphthoquinones are known to form adducts with glutathione even under physiological conditions, and since GSH was the substrate of ChaC1 and was in the reaction buffers at high concentrations (2 mM) this seemed very plausible. We, therefore, examined the inhibitory potential of two juglone-glutathione conjugates. In the absence of purified conjugates to evaluate them experimentally, we examined this using MD simulations. Our results indicated very strong binding of two different conjugates (formed at position 2 and position 3 of the juglone molecule). Thus, it appears that in the presence of ChaC1 when glutathione and juglone form an adduct, this adduct would be inhibitory to the enzyme. Whether the conjugate needs to be formed *in situ* to be effective, or whether it would be effective also when added as a conjugate it is difficult to conclude. However, when we evaluated two other pre-formed conjugates of glutathione, S-methyl glutathione and glutathionyl glutathione, they were both ineffective as inhibitors. Confirmation of this hypothesis of juglone-glutathione conjugates being the inhibitory molecules will thus have to await the availability of these conjugates. However, despite this, it is clear that the inhibitory action of juglone on ChaC1 is novel, since juglone does not appear to be inhibiting ChaC1 by any of the known and reported mechanisms.

Both the two yeast screens that we developed were very robust and amenable to adaptation to a high throughput robotics system. Of these two screens, one of them where the inhibitor allowed growth of the yeast cells was particularly effective. Indeed, the identification of juglone using this screen for an enzyme that has a high substrate Km from a relatively small library of just over 2000 compounds suggests that using this against larger, more complex libraries might yield additional hits against this important target enzyme.

In conclusion, the discovery of juglone and the structurally related plumbagin represent the first leads discovered for inhibition of the ChaC class of enzymes using a robust high throughput screen. The development of these screens and the discovery of these inhibitors is a step forward towards the goal of enhancing glutathione levels in cells by inhibiting glutathione degradation. The current approaches to increasing cytoplasmic glutathione levels are otherwise restricted to increasing biosynthesis (by activating NRF2), or by increasing the supply of glutathione precursors (such as by N-acetyl cysteine) (12,13). Thus, the findings made in this study can allow one to consider a new approach towards this goal, although these compounds may need to be further modified and improved to be effective for use in live cells or organisms.

## Author Contributions

SS initiated the study, performed all the wet lab experiments, and analyzed the results. CC performed and analyzed the *in silico* experiments. DK carried out the molecular dynamic simulations. AKB analyzed the results and supervised the study. SS, CC, and AKB contributed to the writing and preparation of the manuscript. All authors have read and agreed to the manuscript.

## Acknowledgments

We are grateful to Dr. Deepak Sharma and his lab members at the Institute of Microbial Technology, Chandigarh for providing the Spectrum library. We thank Ms. Jasmine for her help with aliquoting the spectrum library. We thank the Robotics facility at IISER Mohali and Mr. Satpal Chawla for their help with Tecan Liquid Handling system programming. We are grateful to Dr Ravi Manjithaya, JNCASR, Bengaluru for his suggestions regarding the high throughput screens and assays. We gratefully acknowledge Dr. SSV Rama Sastry and his student Jay Prakash Maurya of the Department of Chemistry, IISER Mohali for their critical inputs and for providing the juglone analogues and derivatives. SS acknowledges a fellowship from IISER Mohali.

## Conflict of interests

The authors declare that they have no known competing financial interests or personal relationships that could have appeared to influence the work reported in this paper.

## Abbreviations

GSH: glutathione
MD: Molecular Docking
MMGBSA: Molecular mechanics/generalized Born surface area
XP: Extra Precision
SP: Standard Precision
DMSO: Dimethyl sulfoxide
HTS: High Throughput Screening

## Notes

### Competing Interest Statement

The authors have declared no competing interest.

