## Supporting information for "Identification of Juglone, a ‘first-in-class’ inhibitor of the human glutathione degrading enzyme, ChaC1, using yeast-based high throughput screens"

Table S1

| **Name** | **Sequence (5’to 3’)** |
| --- | --- |
| hChaC1NdeI FP | ATGCGCATATGATGAAGCAGGAGTCTGCAGCC |
| hChaC16XHisXhoI RP | CGCATCTCGAGTTAGTGGTGATGGTGATGATGCACCAGCGCCAGAGCCTGCTCGGTGG |
| LEU2 pdr5 FP | ATGCCCGAGGCCAAGCTTAACAATAACGTCAACGACGTTACAACTGTGGGAATACTCAGG |
| LEU2 pdr5 RP | TTATTTCTTGGAGAGTTTACCGTTCTTTTTAGGCACTCTTGCCCTCCTTTTTCTCCTTCTTG |

**Table S2** Details of the Preliminary Hits from Screen I

| **Molecular Name** | **OD_600 nm_** | **Formula** | **M.W** | **Bioactivity** |
| --- | --- | --- | --- | --- |
| CIANIDANOL | 1.00 | C_15_H_14_O_6_ | 290.3 | procollagen production inhibitor, hepatoprotection |
| JUGLONE | 0.66 | C_10_H_6_O_3_ | 174.2 | antineoplastic, antifungal |
| DEOXYCHOLIC ACID | 0.89 | C_24_H_40_O_4_ | 392.6 | bile acid |
| CARBENOXOLONE SODIUM | 0.58 | C_34_H_48_Na_2_O_7_ | 614.7 | anti-inflammatory, antisecretory, antiulcer |
| GOSSYPIN | 0.77 | C_21_H_20_O_13_ | 480.4 | Flavonoid, anti-cancer |
| GLUTATHIONE | 0.927 | C_10_H_17_N_3_O_6_S | 307.3 | antioxidant |
| OXIGLUTATIONE DISODIUM SALT | 0.92 | C_20_H_30_N_6_Na_2_O_12_S_2_ | 656.6 | antioxidant |

OD_600 nm_ refers to the OD_600 nm_ reached by the cultures in the presence of these compounds

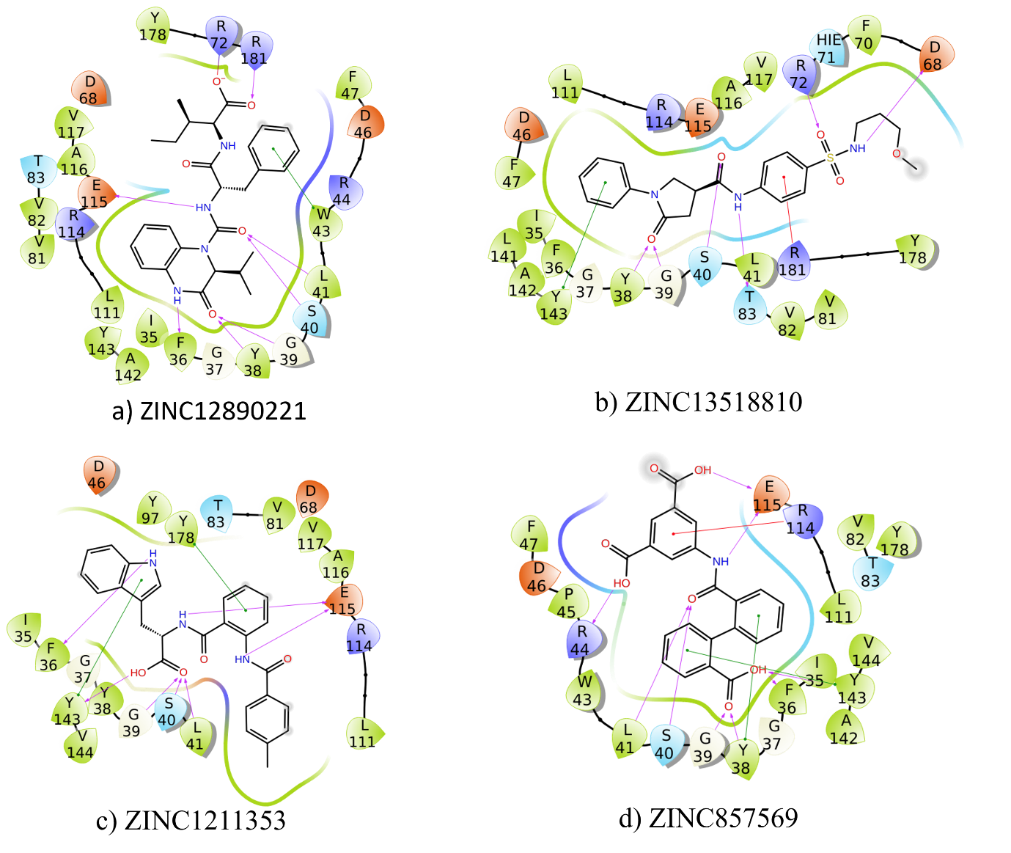

**Figure S1 Protein-ligand interaction patterns obtained after virtual screening** for a) ZINC12890221 b) ZINC13518810 c) ZINC1211353 d) ZINC857569

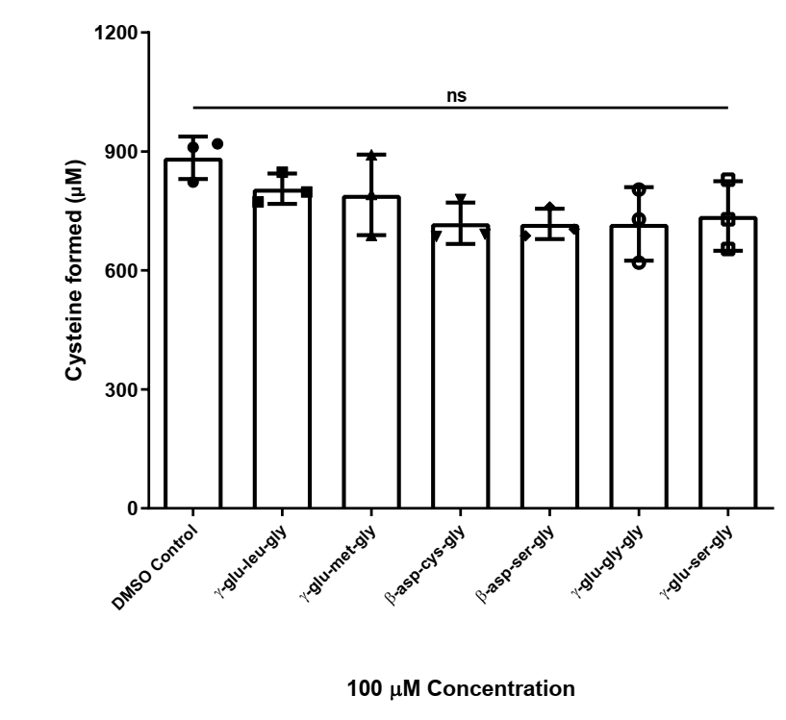

**Figure S2 Effect of glutathione analogues on ChaC1 activity** All glutathione analogues were evaluated at a concentration of 100 μM. The ChaC1p-Dug1p coupled assay was used to estimate the cysteine released as described in methods. The experiment was done twice, with three technical replicates for each sample. The graph here corresponds to the representative data set plotted using the average of the three technical replicates along with ± S.D. The p-value was determined using one-way ANOVA with multiple comparisons. ns: non-significant, p >0.05, * p <0.05, ** p <0.01, *** p <0.001, **** p <0.0001

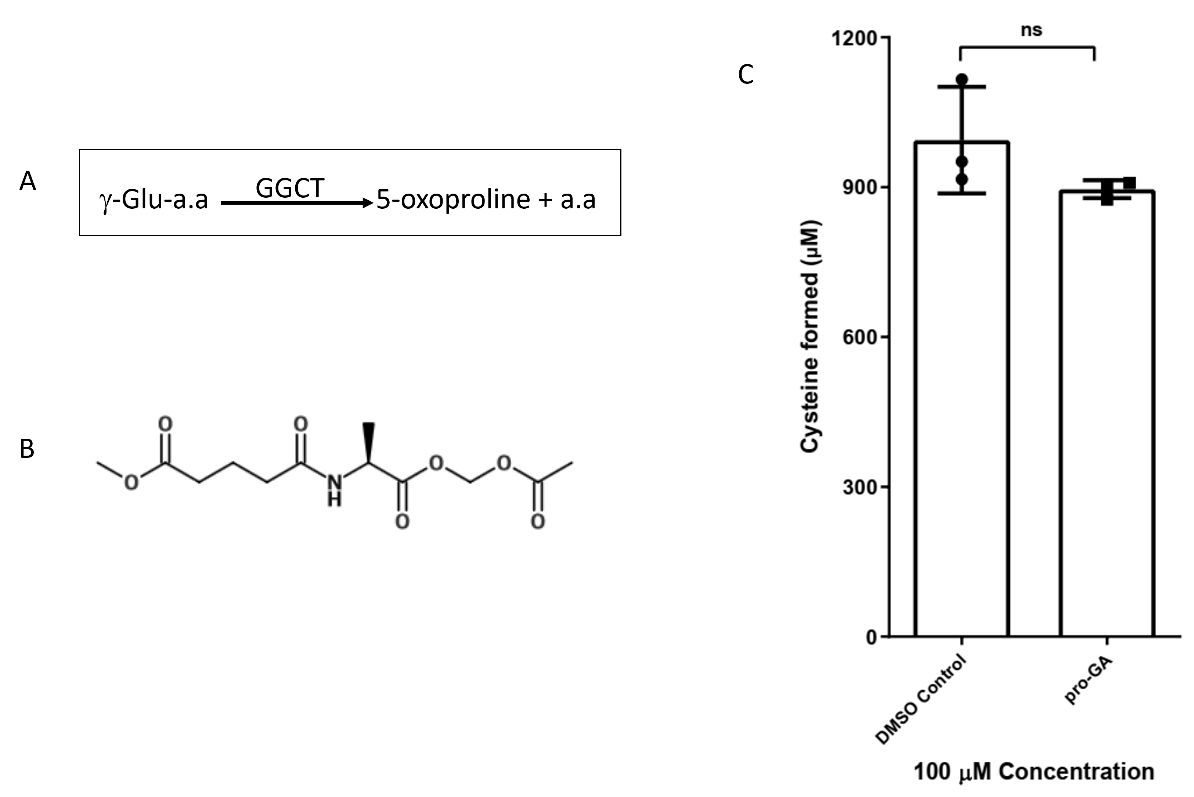

**Figure S3 Effect of pro-GA on ChaC1 activity A Schematic representation of the chemical reaction of GGCT on γ-glu-a.a B Chemical structure of N-glutaryl-L-alanine (GA) C *In vitro* enzymatic assay for pro-GA** pro-GA was dissolved in DMSO and evaluated for its inhibition against the ChaC1 protein at 100 μM concentration with 2 mM of substrate glutathione. The ChaC1p-Dug1p coupled assay was used to estimate the cysteine released as described in methods. The experiment was done once. The graph here corresponds to the representative data set plotted using the average of the three technical replicates along with ± S.D, ns indicates non-significant.

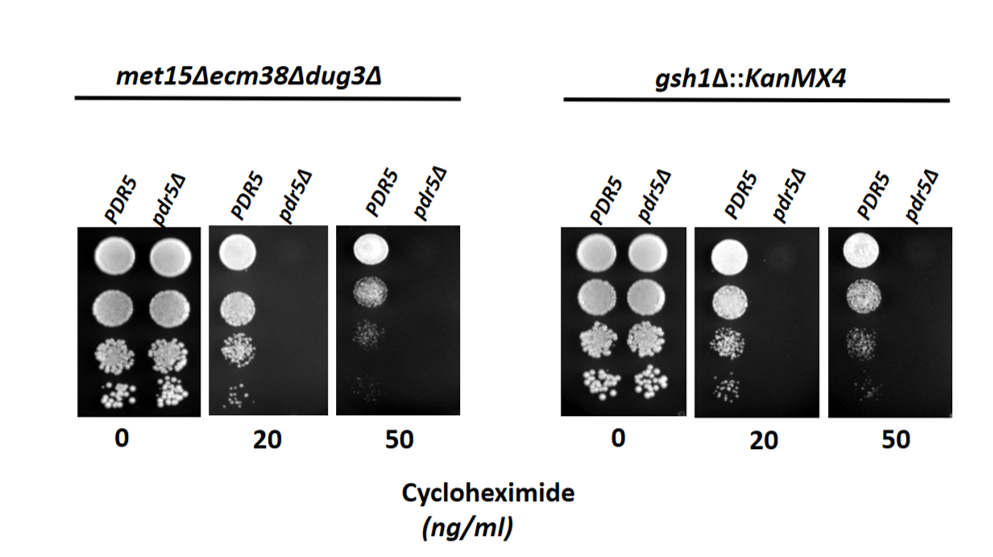

**Figure S4 PDR5 deletion in the yeast strains as seen by their increased sensitivity cycloheximide treatment** *S. cerevisiae* strains ABC1723, ABC1976 transformed with *pdr5∆:: LEU2* deletion cassette (ABC6293 and ABC6303) were evaluated for cycloheximide sensitivity. Strains were grown to exponential phase in a minimal medium, harvested, washed, resuspended in water, and serially diluted to give 0.1, 0.01, 0.001, and 0.0001 OD_600 nm_ of cells. Ten microliters of these dilutions were spotted on SD medium plates containing 0, 20, and 50 ng/mL of cycloheximide. The photographs were taken after 48 hours of incubation at 30℃.

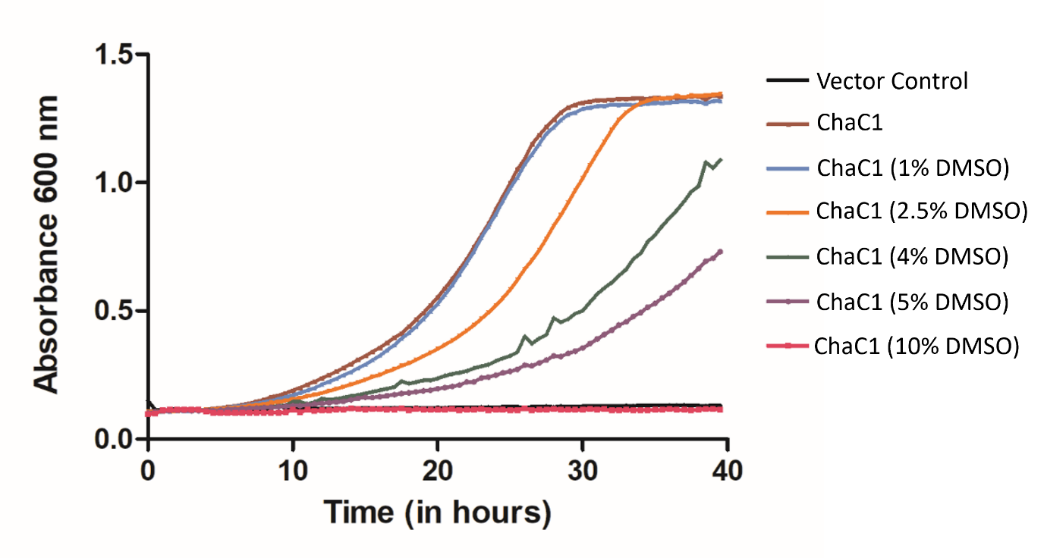

**Figure S5 Screen II analyzed for DMSO toxicity** *S. cerevisiae* strain ABC6293 transformed with ChaC1_WT and the corresponding vector control and was grown overnight in minimal media with methionine as sulfur source. These were then reinoculated at 0.15 OD_600 nm_ in fresh SD medium without any sulfur source and grown until the cells reached the early exponential phase. The yeast cells were again reinoculated at 0.15 OD_600 nm_ in fresh SD medium with 100 μM GSH along with 0, 1, 2.5, 4, 5, and 10% DMSO concentrations in 96-well microtiter plates. Their growth was monitored for 40 hours. The experiment was done once. The graph here corresponds to the representative data set plotted using the average of three technical replicates along with ± S.D. values.

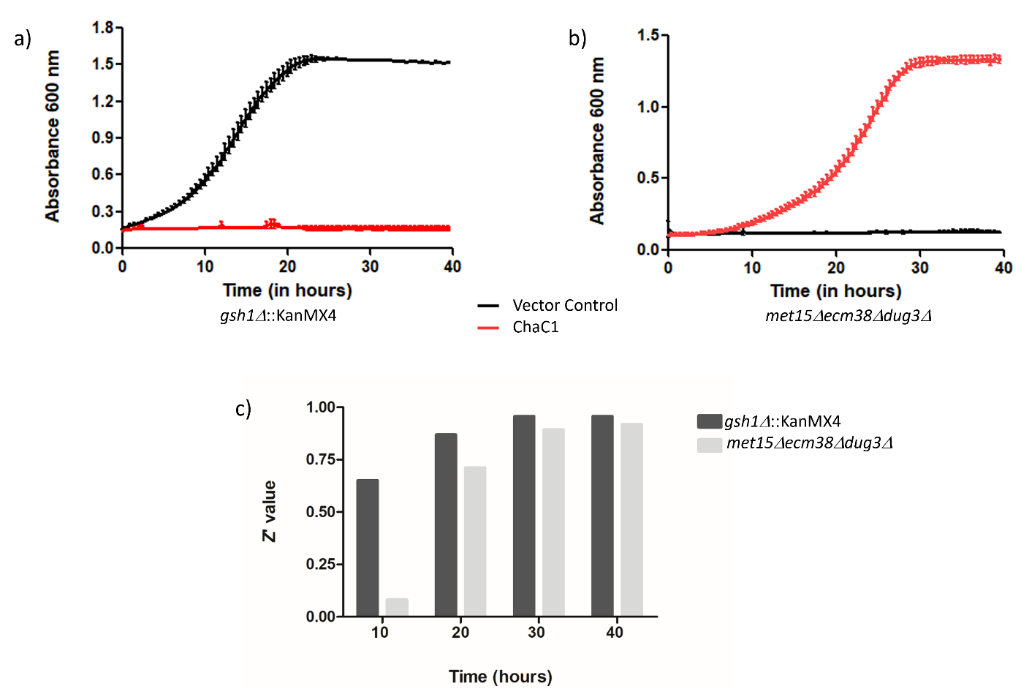

**Figure S6 Growth curve analysis for Screen I and Screen II** a) and b) *S. cerevisiae* strains ABC 6293 and ABC 6303were transformed with ChaC1_WT and the corresponding vector control and were grown overnight in minimal media with glutathione (for Screen I) and methionine (for Screen II) as sulfur sources. These were then reinoculated at 0.15 OD_600 nm_ in fresh SD medium without any sulfur source and grown until the cells reached the early exponential phase. The yeast cells were again reinoculated at 0.15 OD_600 nm_ in fresh SD medium without any sulfur source (Screen I) and 100 μM GSH (Screen II) in 96-well microtitier plates. Their growth was monitored for 40 hours. Three datasets for each sample from were obtained and plotted using graph pad prism software. c) Z’ values for both the screens approach ~0.9 over 40 hours. The experiment was done multiple times, along with three technical replicates for each sample. The graph here corresponds to the representative data set plotted using the average of the three technical replicates along with ± S.D. values.

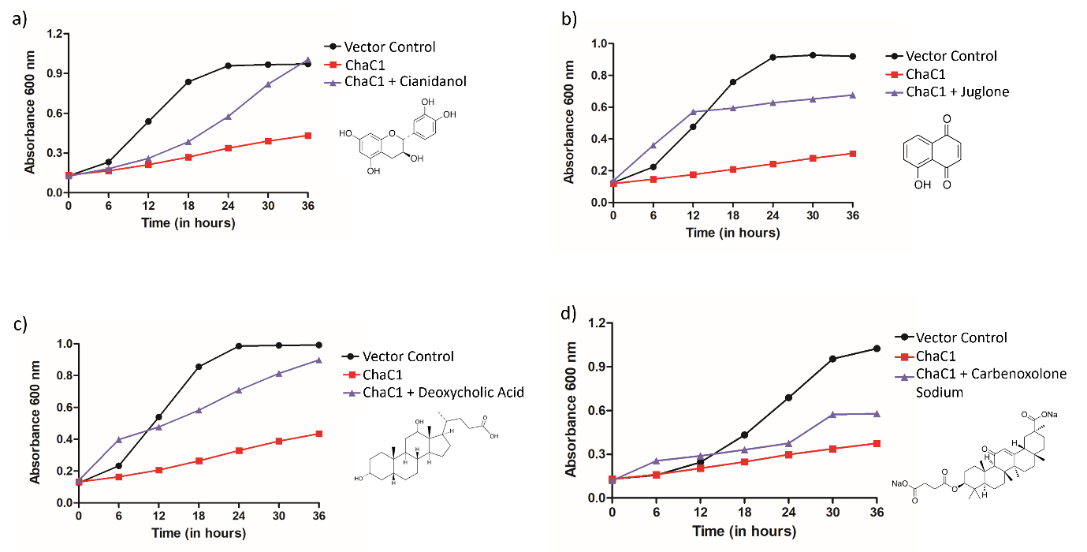

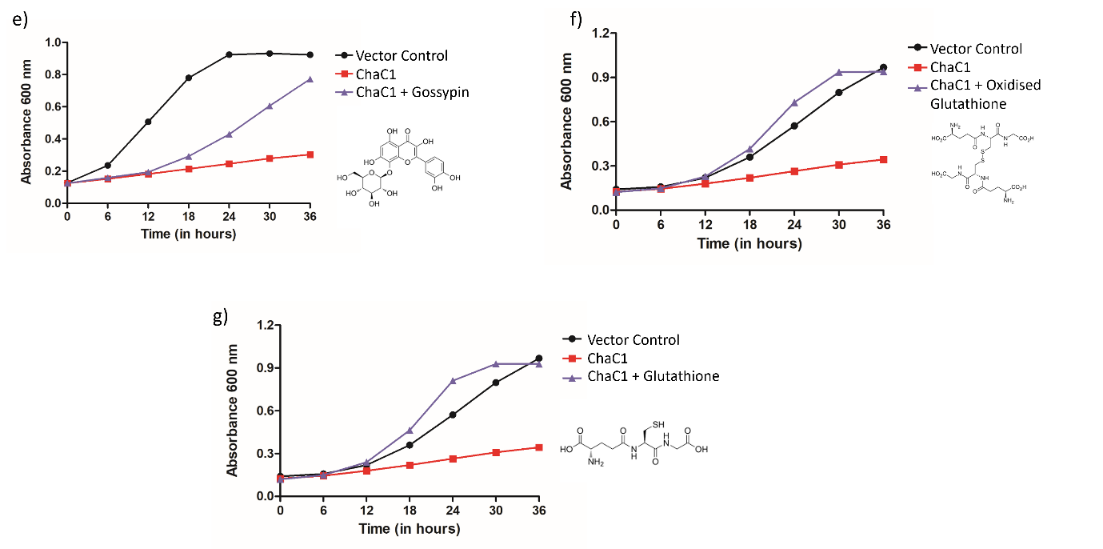

**Figure S7 Individual growth curves plotted for the hits at 100 μM concentrations** a) Cianidanol b) Juglone c) Deoxycholic acid d) Carbenoxolone sodium e) Gossypin f) Oxidized Glutathione g) Glutathione

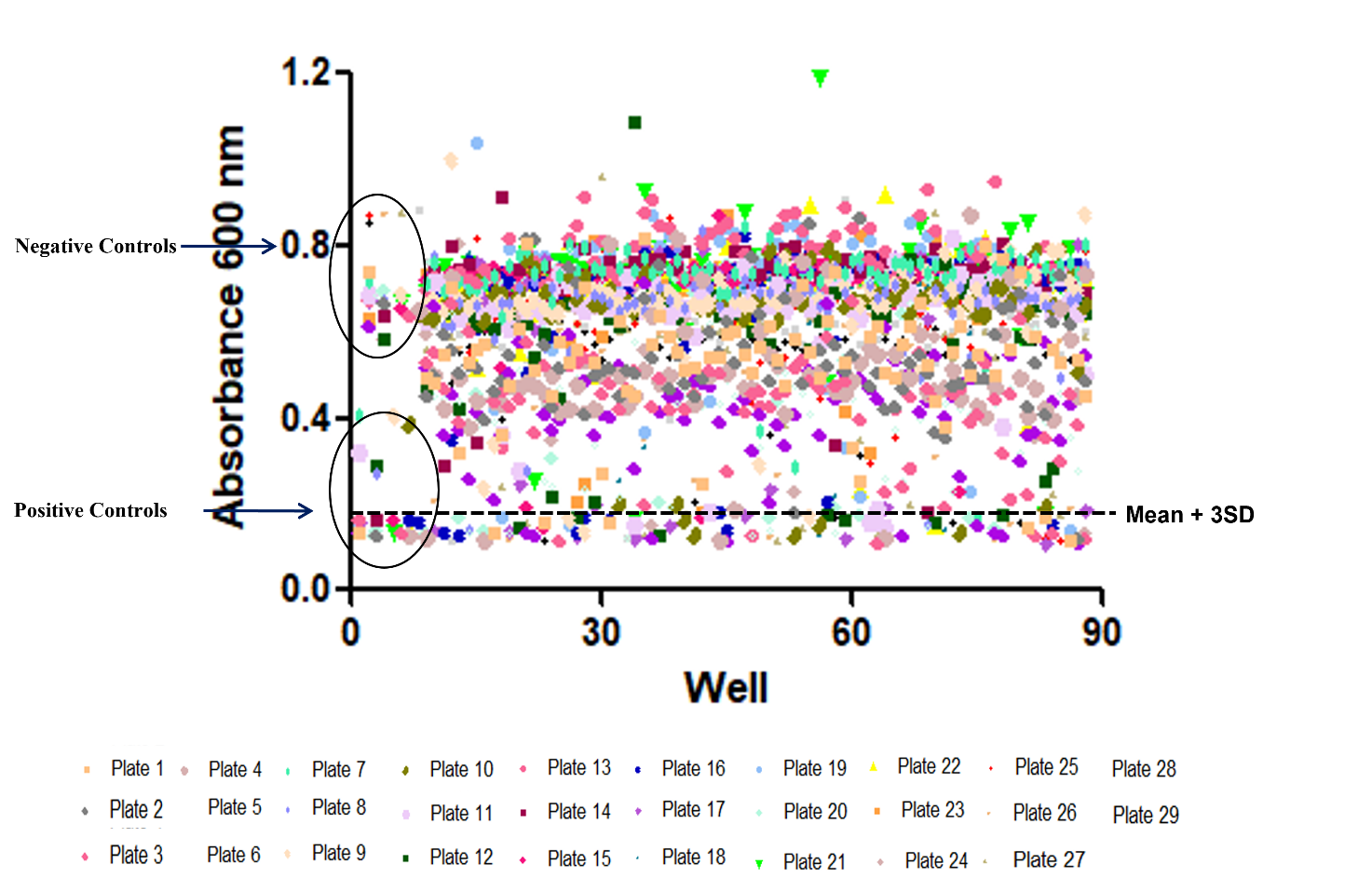

**Figure 5.7 High throughput screening of Spectrum library using Screen II (sulfur auxotrophic yeast strain)** *S. cerevisiae* strain ABC 6293 transformed with ChaC1 and the corresponding vector control was grown according to the growth conditions optimized for High-throughput screening assay (Methods Section 2.2.16). 200 μL (0.15 OD_600 nm_) of ChaC1 expressing yeast cell culture was dispensed in six 96-well microtiter plates and 100 μM (10 μL) of each compound from the Spectrum library was added using a liquid handling system. Controls with only DMSO were also added to each plate. The plates were incubated at 30℃ with continuous shaking in the incubator and the growth was monitored for 36 hours. Endpoint A_600_ was obtained for each compound and plotted using graph pad prism software. The dotted line indicates three standard deviations above the mean effect.

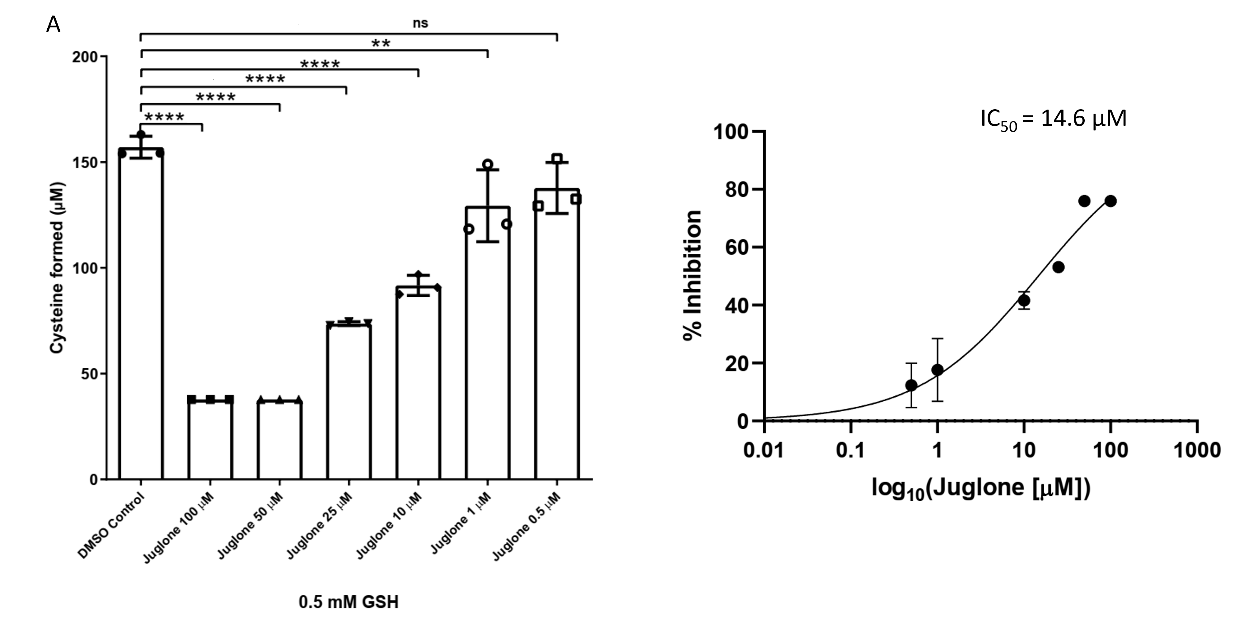

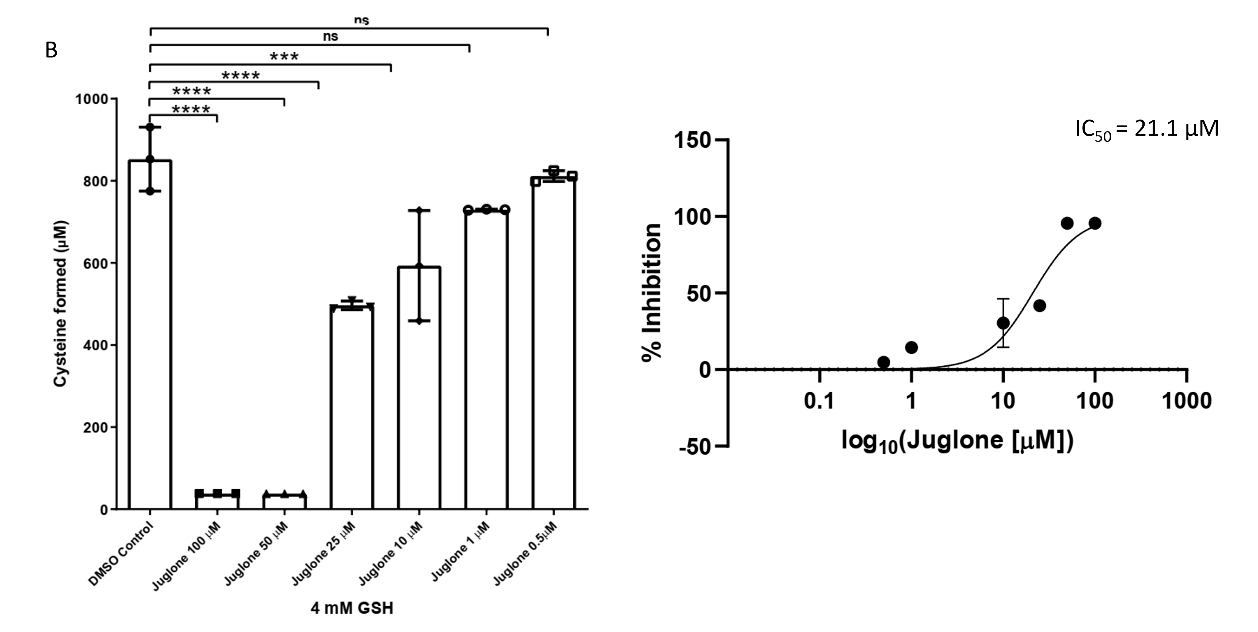

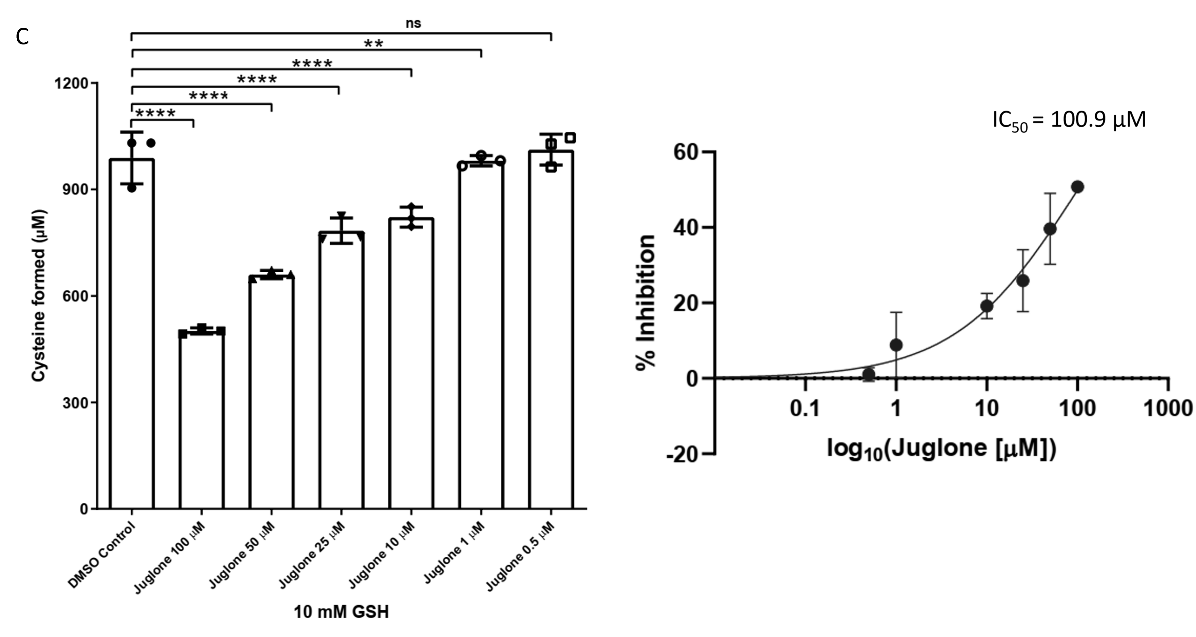

**Figure S9 Determination of IC_50_ of juglone against ChaC1 enzyme** Different concentrations of juglone were evaluated for their inhibition against the ChaC1 protein. The ChaC1p-Dug1p coupled assay was used to estimate the cysteine released as described in methods. The IC_50_ value for juglone was determined against glutathione concentrations of A 0.5 mM GSH B 4 mM GSH and C 10 mM GSH. The experiment was done thrice, along with three technical replicates for each sample. The graphs here correspond to the representative data set plotted using the average of the three technical replicates along with ± S.D. values. The p-value was determined using one-way ANOVA with multiple comparisons. ns: non-significant, p>0.05, *p<0.05, **p<0.01, ***p<0.001, ****p<0.0001

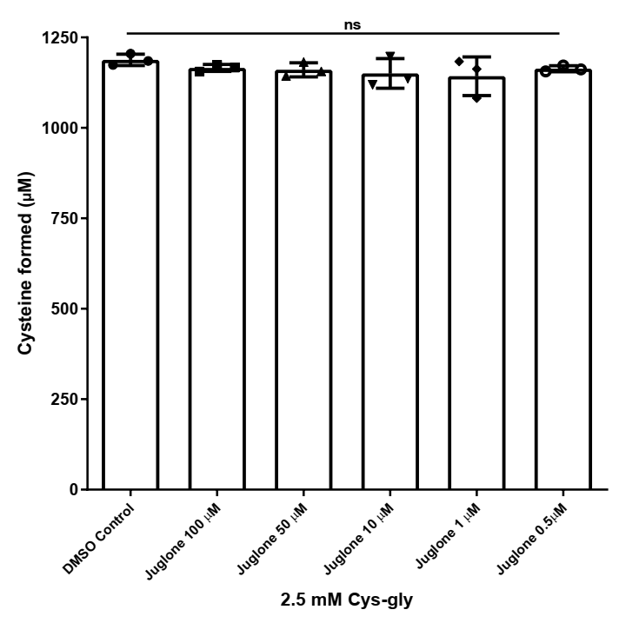

**Figure S10 Evaluation of the inhibitory effect of juglone on Dug1 protein** Juglone concentrations ranging from 0.5 μM to 100 μM were evaluated for their inhibition against the Dug1 protein. Acidic ninhydrin assay was used to estimate the cysteine released as described in the methods. The experiment was once. The graph here corresponds to the representative data set plotted using the average of the three technical replicates along with ± S.D. The p-value was determined using one-way ANOVA with multiple comparisons. ns: non-significant

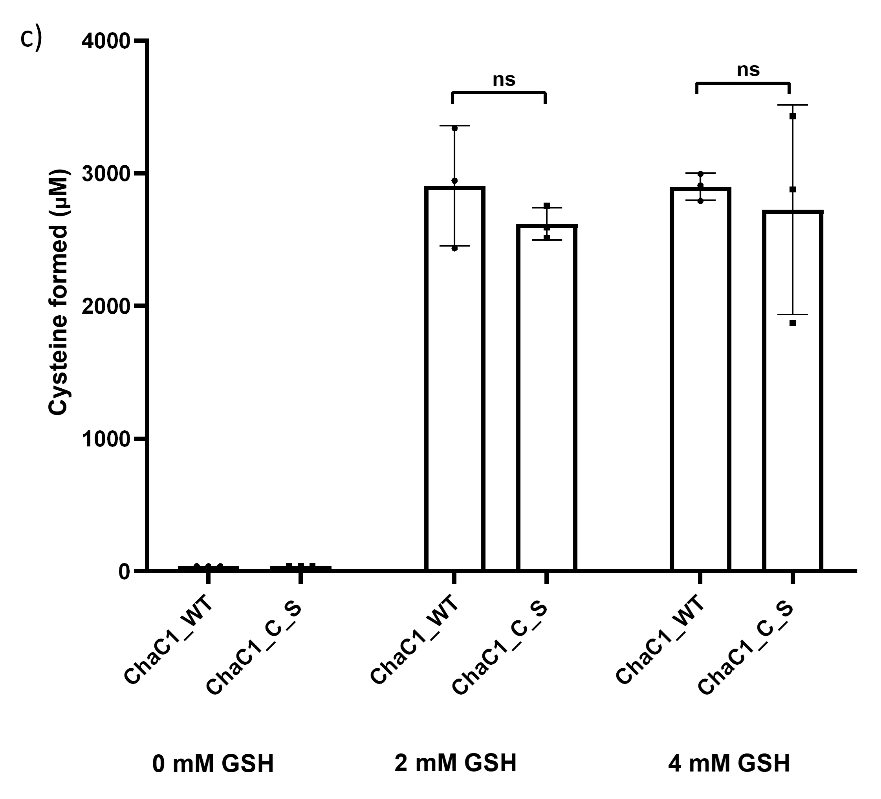

**Figure S11 Relative activity of ChaC1_C_S mutant** 0 mM, 2 mM and 4 mM concentrations of glutathione were used to determine the specific activities for ChaC1_WT and ChaC1_C_S mutant. The ChaC1p-Dug1p coupled assay was used to estimate the cysteine released as described in the methods. The experiment was done thrice, along with three technical replicates for each sample. The graph here corresponds to the representative data set plotted using the average of the three technical replicates along with ± S.D. values.

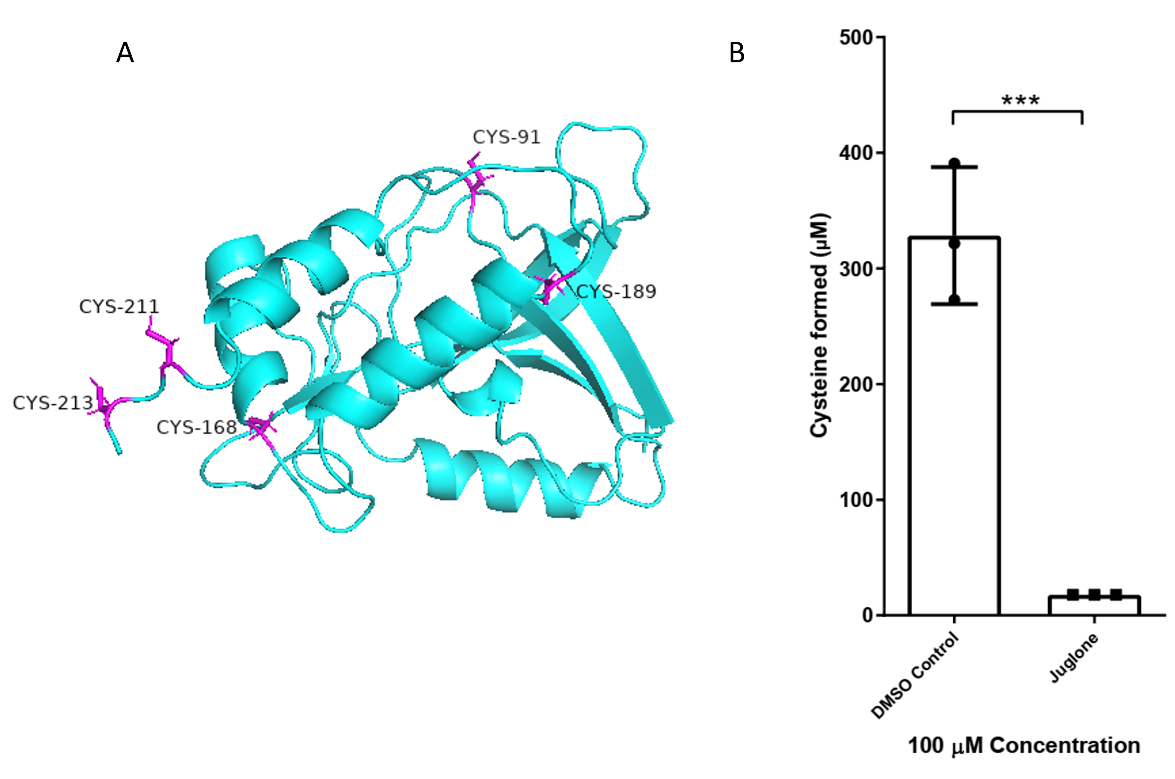

**Figure S12 Inhibition of juglone against the Cysteine-free ChaC1 mutant A 3-D representation of cysteine-free ChaC1 mutant** The figure highlights (in dark pink) Cys91, Cys168, Cys189, Cys211 and Cys213 that have been mutated to serine in the ChaC1 model (cyan colour) generated by homology modelling **B Effect of juglone on ChaC1_C_S mutant** Juglone was evaluated for its inhibition against the ChaC1_C_S protein at 100 μM concentration with 2 mM of substrate glutathione. The ChaC1p-Dug1p coupled assay was used to estimate the cysteine released as described in methods. The experiment was done twice, along with three technical replicates for each sample. The graph here corresponds to the representative data set plotted using the average of the three technical replicates along with ± S.D. values. The p-value was determined using unpaired t-test. ns: non-significant, p>0.05, *p<0.05, **p<0.01, ***p<0.001, ****p<0.0001

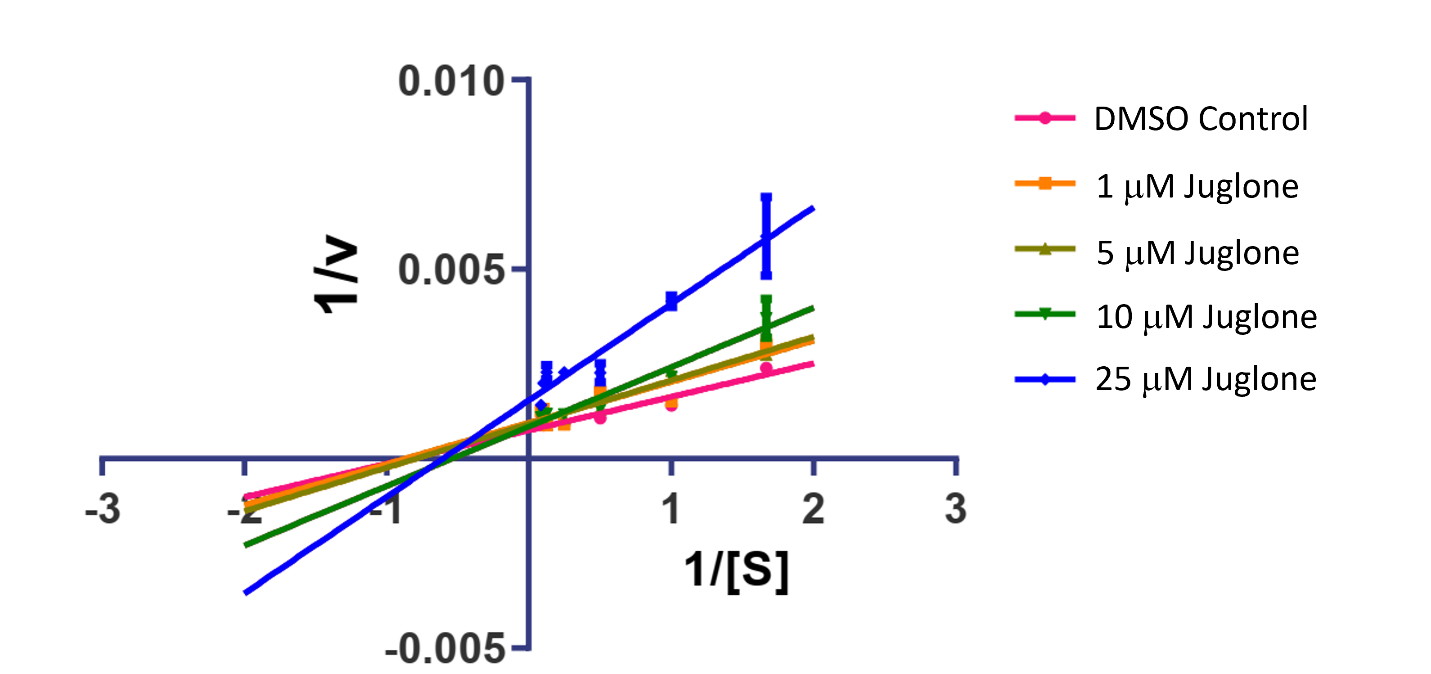

**Figure S13 Lineweaver-Burk plot showing mixed inhibition of ChaC1 by juglone** Different concentrations of juglone were evaluated against varying concentrations of the substrate, glutathione (0.6 mM, 1 mM, 2 mM, 4 mM, 8 mM, 10 mM, and 12 mM). The ChaC1p-Dug1p coupled assay was used to estimate the cysteine released as described in methods. The graph here corresponds to the representative data set plotted using the average of three technical replicates along with ± S.D. values.

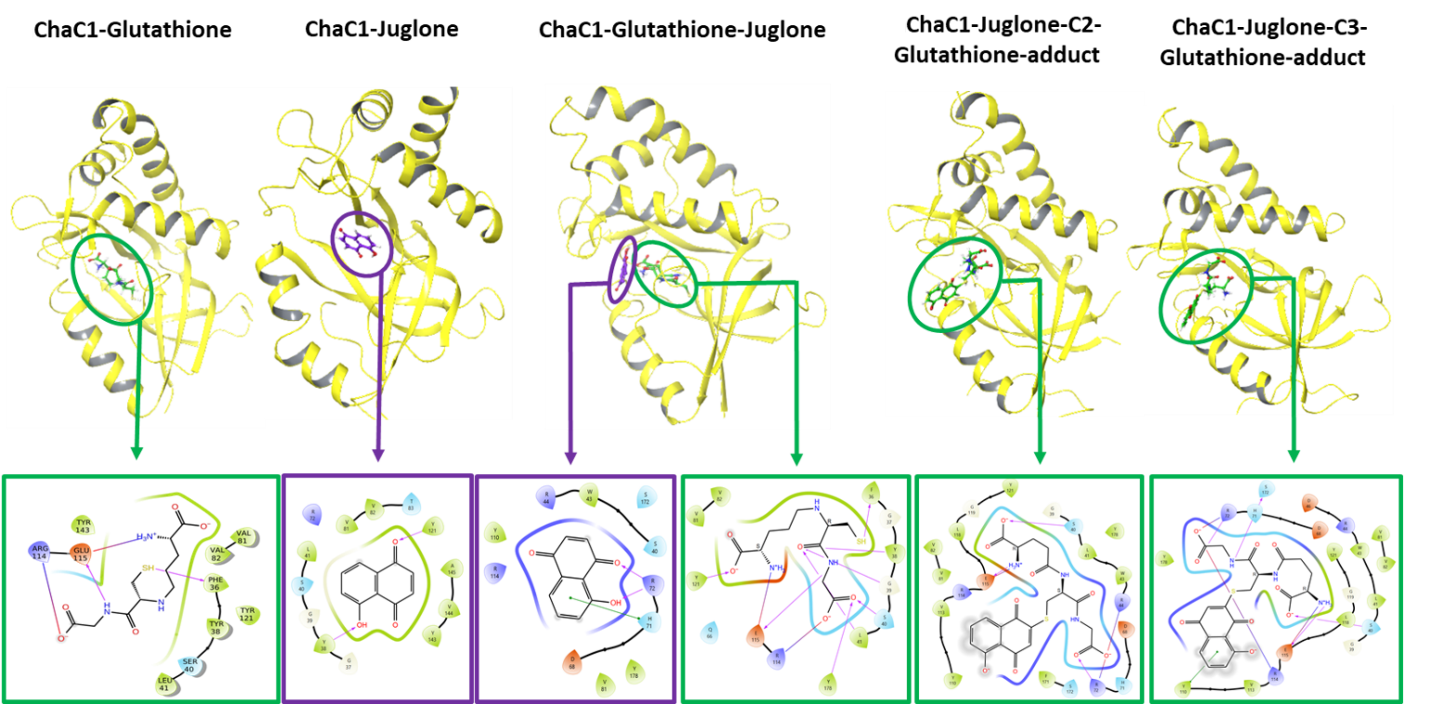

**Figure S14** Initial binding positions and interactions of the ligands (glutathione, juglone, and both the adducts) with ChaC1 before MD simulations.
